# Origin tracing and detection lag in the first *Aedes albopictus* invasion of mainland Australia and a long-term genomic dataset of mosquito incursions

**DOI:** 10.64898/2026.11.19.767973

**Authors:** Thomas L Schmidt, Odwell M Muzari, Nancy M Endersby-Harshman, Christopher Paton, Joe Davis, Gerhard Ehlers

## Abstract

Proactive management of invasive species aims to prevent and eradicate new incursions before they establish. Genomic data can assist proactive management around how to deploy resources and when to change strategies, and has been used effectively in managing *Aedes albopictus* (Asian tiger mosquito) in the Australian Northern Peninsula Area over the past decade. This study characterises the first major incursion of *Ae. albopictus* on the Australian mainland, and contrasts this with long-term genomic monitoring of this species within the Torres Strait Islands (TSI). First, we genomically traced the mainland incursion, which was detected in Cape York, Queensland, to the nearest infested island, Muralug (Prince of Wales). Strong genetic drift patterns in the incursion are suggestive of recent inbreeding, and point to a lag between incursion and detection of at least two mosquito generations. This stands in stark contrast with incursions within the TSI, which showed no evidence of inbreeding, suggesting no detection lag in the TSI. We genomically traced n=63 incursions detected in the TSI between 2019–2026, including n=44 newly sequenced samples; this used an expanded reference databank containing samples from one island, Muralug, that had not been available in previous tracing. Including Muralug greatly improved tracing success from ∼50% to 77.8—92.1%. We observed no reduction in tracing success when whole genome and reduced representation sequencing datasets were combined in analysis, indicating composite approaches are viable. Together these findings demonstrate the value of genomic databanks for long-term incursion monitoring, provided these capture a sufficient proportion of genetic variation.

## Introduction

The global increase in human movement and transportation has led to attendant risks in the accidental transport of other organisms. New populations that establish as a result of this transport are termed biological invasions, and these can confer costs to the local environment or to the health of humans, livestock, and crops (Bradshaw et al., 2016). Before an invasive population becomes established, it first has to disperse to the new environment (i.e. an incursion), after which it must persist for a number of generations while it grows to sufficient size that eradication becomes impracticable (i.e. an established invasion). This initial period, between incursion and establishment, is critical for biosecurity, as it defines the window of time in which the population can be detected and eradicated.

For areas of high biosecurity risk and with entrenched biosecurity protocols, proactive management can be used to detect incursions and eradicate them before they become established (Ahmed et al., 2022; Cuthbert et al., 2022). In such cases, evolutionary assessment of specific incursions can provide insight into where incursions are coming from, the number of independent incursions, and the time lag between dispersal and detection. Origins can be inferred through population assignment (Chen et al., 2018) or locational inference (Battey et al., 2020), and can help identify locations of differential biosecurity risk, allowing resources to be deployed more efficiently (Bacon et al., 2012; Caley et al., 2015). For incursions detected simultaneously, their independence can be assessed through pairwise relatedness, where individuals from the same kinship group are likely part of the same rather than independent incursions; this can also point to differential biosecurity risk. Detection lag can be measured by how many generations have been produced in the new environment, which in populations that usually outbreed can be inferred through patterns of founder effects in dispersers (F0 generation), genetic drift (F1 generation onwards), and inbreeding (F2 generation onwards). Detection lag indicates how effective current biosecurity protocols are at detecting new incursions after they arrive. If incursion monitoring is conducted over multiple years, these become valuable datasets for long-term monitoring (Stroud C Ratcliff, 2025), where they can help assess how risks change gradually or in response to sudden environmental changes (Lassaline et al., 2026).

The Torres Strait Islands (TSI) in northeastern Australia represent an area of high biosecurity risk. The broader geographical region comprises southern Papua New Guinea (PNG), the inner and outer Torres Strait Islands, and Cape York Northern Peninsula Area on the Australian mainland (Fig. 1a). Many biological invasions have spread into and through this region (Ridland et al., 2025; Ritchie et al., 2006; Waltham et al., 2023; Xu et al., 2022). One invasion of particular significance is *Aedes albopictus*, the Asian tiger mosquito, which has displaced *Ae. aegypti* from many islands of the Torres Strait (Muzari et al., 2019) where it is now the primary vector for dengue (Muzari et al., 2026). *Aedes albopictus* established on some of the outer islands in 2004, and spread rapidly to other islands thereafter, reaching the inner islands of Ngurapai (Horn) and Waiben (Thursday) in 2010 (Muzari et al., 2017). The species became densely established on Ngurapai and Waiben but was eradicated from the two islands between 2011 and 2015 through an intensive vector control program (Muzari et al., 2017). Since this time, a holding pattern using the *cordon sanitaire* biosecurity approach has been maintained (van den Hurk et al., 2016), with *Ae. albopictus* remaining abundant throughout all islands apart from Ngurapai and Waiben, and with no invasions detected on the Australian mainland including the Cape York region until 2026.

**Figure 1.**
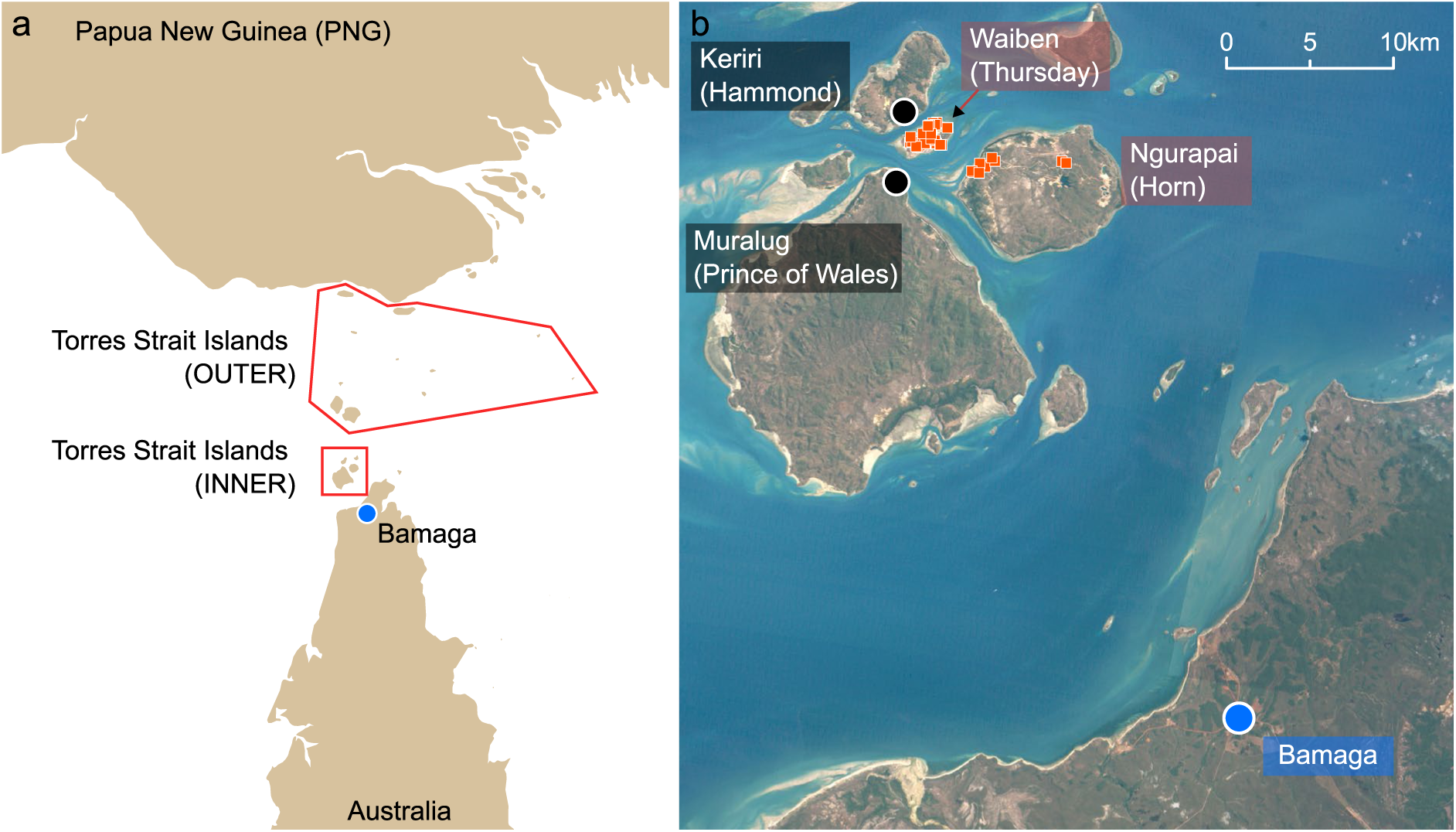
Sample site maps. (a) Shows mainland Australia and PNG, the outer Torres Strait Islands (all with *Ae. albopictus*), and the inner Torres Strait Islands (some with and some without *Ae. albopictus*). Blue circle shows location of mainland incursion of Australia in Bamaga, Cape York. (b) The inner Torres Strait Islands. Keriri and Muralug both have *Ae. albopictus*, the black circle shows the location of villages where *Ae. albopictus* were sampled for the reference databank. Waiben and Ngurapai are free from *Ae*. *albopictus* since about 2015, but have experienced occasional *Ae. albopictus* incursions, with analysed incursions (2019-2026) displayed as red squares. Each incursion was successfully eradicated soon after detection. Maps produced in ArcGIS Pro (version 3.7); (b) uses the Torres Strait Clear sky Landsat (https://eatlas.org.au/data/ uuid/71c8380e-4cdc-4544-98b6-8a5c328930ad).

Occasional isolated incursions of *Ae. albopictus* have been detected on Ngurapai and Waiben after the successful eradication. Beginning in 2019, evolutionary genomics has been used to infer the origins of these incursions via a genomic reference databank (Schmidt et al., 2021, 2023). These analyses show consistent patterns, where most incursions are from nearby islands, but with occasional long-distance movement from farther afield. There has been no evidence of inbreeding or strong genetic drift suggestive of cryptic establishment, indicating that current biosecurity protocols have been effective at detecting incursions before they establish (Schmidt et al., 2023). The genetic profile of TSI *Ae. albopictus* is derived from its invasion source in or around Indonesia (Beebe et al., 2013; Maynard et al., 2017) but with additional admixture with PNG genetic backgrounds (Schmidt et al., 2021). This admixture, along with the drift associated with invasion bottlenecks, makes it possible to trace mosquitoes to individual islands within the TSI (Schmidt et al., 2021, 2023). Insecticidal approaches have been sufficient to eradicate *Ae. albopictus* incursions and reduce populations on invaded islands during dengue outbreaks (Muzari et al., 2017). Alleles conferring target-site resistance to pyrethroids are apparently absent from TSI *Ae. albopictus* but have been recorded in nearby populations in PNG, Timor-Leste, and Vanuatu (Schmidt et al., 2024).

In April 2026, an incursion of *Ae. albopictus* was detected on the Australian mainland for the first time in Bamaga, Queensland, (Fig. 1) in the Cape York Northern Peninsula Area. This led to an immediate response involving further surveillance, residual harbourage spraying on fringing vegetation, and yard-to-yard treatment of larval habitats throughout the Bamaga community. The *Ae. albopictus* specimens that were detected at this point were also sampled for subsequent evolutionary genomic analysis. This study presents the results of this analysis, aiming to infer the origin of the incursion and estimate how much time had passed between incursion and detection (detection lag). Results are interpreted with reference to those from other incursions detected on Ngurapai and Waiben between 2019–2026. Finally, methods for generating genomic data have changed in cost since 2018, when the first data were produced for this project. Accordingly, the TSI genomic tracing program has changed its workflows for new incursions from double digest RAD sequencing (Peterson et al., 2012) to whole genome sequencing. We evaluate whether analysing these two data types together may negatively affect tracing success, and also evaluate whether tracing success would be improved when an additional island was added to the reference databank (Muralug (Prince of Wales Island), Fig. 1b).

## Methods

### New samples of Aedes albopictus

#### Mainland incursion

In April 2026, vector surveys were conducted in Bamaga (Fig. 1b) and four other Cape York communities of Seisia, New Mapoon, Umagico and Injinoo, aiming to detect possible incursions of *Ae. albopictus* onto the Australian mainland. Sampling followed the sweep-net methods used previously (Muzari et al., 2017), and focused on selected sites located in suitable *Ae. albopictus* sylvatic habitat around and within communities. Bamaga had detections of *Ae. albopictus*, with nine adults (5 males and 4 females) collected in mid-April, and eight adults (7 males and 1 female) collected about two weeks later in late April. Genomic analysis was conducted on six specimens from mid-April and seven from late April. Similar surveys have been conducted in the same communities annually in the wet season for at least the previous 10 years, with no detection of *Ae. albopictus*.

#### TSI incursions

Between 2023 and 2026, sweep-net surveys for *Ae. albopictus* were conducted on Ngurapai (Horn) and Waiben (Thursday) islands. Surveys were conducted 3-4 times each wet season at selected sites with suitable vegetation across each island. Adult *Ae. albopictus* were detected in 2023, 2025 and 2026. Genomic analysis was conducted on every sample, treating each putatively as an independent incursion. Incursions detected before 2023 were analysed in previous work (Schmidt et al., 2021, 2023).

#### Additional reference samples

Previous tracing work in TSI made use of a genomic reference databank (n=240 mosquitoes) from 12 villages. However, this dataset did not include Muralug (Prince of Wales), an inhabited island near to Waiben and Ngurapai but with an extant *Ae. albopictus* population (Fig. 1b). We reasoned that the absence of Muralug from the dataset could negatively impact tracing results, leading to fewer incursions being traced with accuracy and precision. Accordingly, we sampled *Ae. albopictus* from Muralug to add them the reference databank. Adults were sampled using human-baited sweep nets in April 2023, with samples taken at 100 m separation distance to reduce close kin sampling, following identical protocols as previously-collected reference samples (Schmidt et al., 2021). We selected 20 Muralug *Ae. albopictus* for sequencing.

### DNA sequencing, genotyping, and filtering

#### New and old sequence data

Different methods were used for generating genomic data between 2023 and 2026, reflecting changing costs of sequencing and library preparation. For samples collected in 2023, data were generated using double digest restriction-site associated DNA sequencing (ddRADseq, (Peterson et al., 2012)), following identical methods to those used to produce the reference databank (Schmidt et al., 2021). This included 28 incursion samples (n=27 Waiben; n=1 Ngurapai) and the reference samples from Muralug (n=20). For samples collected in 2025–2026, data were generated using whole genome sequencing (WGS). This included 16 incursion samples (n=7 Waiben 2025; n=1 Waiben 2026; n=8 Ngurapai 2026) and the samples from the mainland incursion (n=13). For 2023 samples processed with ddRADseq, genomic DNA was extracted using Qiagen DNeasy® Blood C Tissue Kits (Qiagen, Hilden, Germany). Extracted DNA was used to build ddRAD libraries following the protocol of Rašić et al. (2014). Libraries were individually barcoded, then sequenced on either a HiSeq 4000 or a Novaseq 6000 using 150 bp chemistry. For 2025–2026 samples processed with WGS, whole mosquitoes were sent to BGI Genomics for DNA extraction, quantification, and sequencing, targeting 20 Gb clean data per sample.

Two reference databanks were used for tracing and other evolutionary analyses. These included a TSI-only databank containing n=257 samples from 13 TSI villages, and a global databank of n=523 samples from 37 populations across 20 countries. All reference data were produced using ddRADseq following the same methods as above and previously published (Schmidt et al., 2021, 2024), except for the n=44 samples from USA, Guam, and Marshall Islands which were previously published sequences produced using WGS (methods described by Seok et al. (2025)). Previously published data are accessible by NCBI sequence read archive (BioProjects PRJNA1118433 and PRJNA1227508).

Finally, to test whether the inclusion of Muralug samples in the reference databank improved tracing analyses, we reanalysed 20 incursion samples from previous years (n=4 Ngurapai 2019; n=3 Ngurapai 2021; n=4 Waiben 2021; n=5 Ngurapai 2022; n=4 Waiben 2022). These were produced using ddRADseq following the same methods as above.

#### Alignment, genotyping, and filtering

All genomic data were aligned to the AalbF5 nuclear genome assembly for *Ae. albopictus* (GenBank: GCA_035046485.1). Alignment was with Bowtie2 v2.3.4.3 (Langmead C Salzberg, 2012) with “–very-sensitive” settings. Following alignment, bam files were filtered with samtools v1.16 to retain only read pairs mapped to primary alignments. For ddRAD samples, an additional demultiplexing step was required before alignment, which was done using the Stacks v2.54 programme “process_radtags” (Rochette et al., 2019). For WGS samples, an additional duplicate removal step was required after alignment, which was done using the Picard programme “MarkDuplicates” (http://broadinstitute.github.io/picard/).

For analysis of incursions from 2023 and earlier, the Stacks programmes “ref_map” and “populations” were used to call SNP genotypes and export VCF files containing SNPs called in at least 50% of individuals from each population and 90% of individuals total, and with minor allele count ≥ 3 (Linck C Battey, 2019). All samples had <25% missing data, and the total missing data was low (x̅ = 5.04%).

For analysis of incursions from 2025–2026, genotyping was with GATK v4.2.6.1 (Van der Auwera C O’Connor, 2020). First, we extracted the SNPs called in the above ddRAD analyses, and designated these as interval sets for genotyping WGS data. This ensured the same SNPs were used in ddRAD and WGS analyses. We then used GATK “HaplotypeCaller” to call haplotypes at these SNPs in reference and incursion samples, with an additional 100 bp padding around each SNP. Samples were genotyped together using “GenomicsDBImport” to import the samples to a database then “GenotypeGVCFs” to call genotypes. We used “SelectVariants” and “VariantFiltration” to remove indels and filter SNPs following standard GATK guidelines for non-model taxa, setting “QD < 2.0”, “QUAL < 30.0”, “SOR > 3.0”, “FS > 60.0”, and “MQ < 40.0”.

Following filtering, missing data were imputed in windows of 50 SNPs using Beagle v4.1 (Browning C Browning, 2016), using 10 iterations, 10 SNP overlaps, and the full dataset as reference.

### Population genomic analysis

#### Mainland incursion

To infer origins, we used PCA, run in R package “LEA” (Frichot C François, 2015), and Treemix v1.13 (Pickrell C Pritchard, 2012). To infer histories of genetic drift, we used Tajima’s D (Tajima, 1989) and pairwise relatedness. Tajima’s D was run in VCFtools (Danecek et al., 2011), using 100 kb windows and omitting windows with fewer than 5 SNPs. Relatedness analysis used the KING approach (Manichaikul et al., 2010) and was run in PLINK v1.90b6.16 (Chang et al., 2015). This approach to relatedness has been shown to produce reliable results for reduced representation datasets under a range of conditions (McMaster et al., 2025).

#### Incursions into the inner Torres Strait Islands

To infer origins, we used Locator (Battey et al., 2020). The reference dataset had already undergone cross-validation in Locator, for which 86% of reference samples were assigned unambiguously to their correct island (Schmidt et al., 2023). Locator was run with 1000 bootstrap replicates to indicate confidence in estimates. To infer the independence of the incursions, we used pairwise relatedness run in PLINK as described above. Inferring exact relationships is difficult in recently invasive systems as these have high residual coancestry that leads to patterns of relatedness even among unrelated individuals (Schmidt et al., 2021). We assigned related pairs using a cut-off of 0.088, which represents 2^nd^ order or higher relatedness (Manichaikul et al., 2010).

## Results

### Mainland incursion

#### Origin of mainland incursion

PCA indicated that the mainland incursion was of a single genetic group (Fig. S1). PCA also plotted the mainland sample apart from TSI samples, which could indicate either a non-TSI origin, or strong genetic drift as expected from an invasion bottleneck (c.f. Schmidt et al. (2023)). We explored this further using Treemix, which explicitly models differential rates of genetic drift among sets of populations. Treemix of the global dataset placed the mainland incursion clearly within the TSI (Fig. 2a). This placement, along with the long internal branch leading to the mainland incursion, suggests that the mainland incursion had an origin in the TSI but has experienced strong genetic drift. Treemix of the TSI-only dataset indicated a likely origin in Muralug (Fig. 2b).

**Figure 2.**
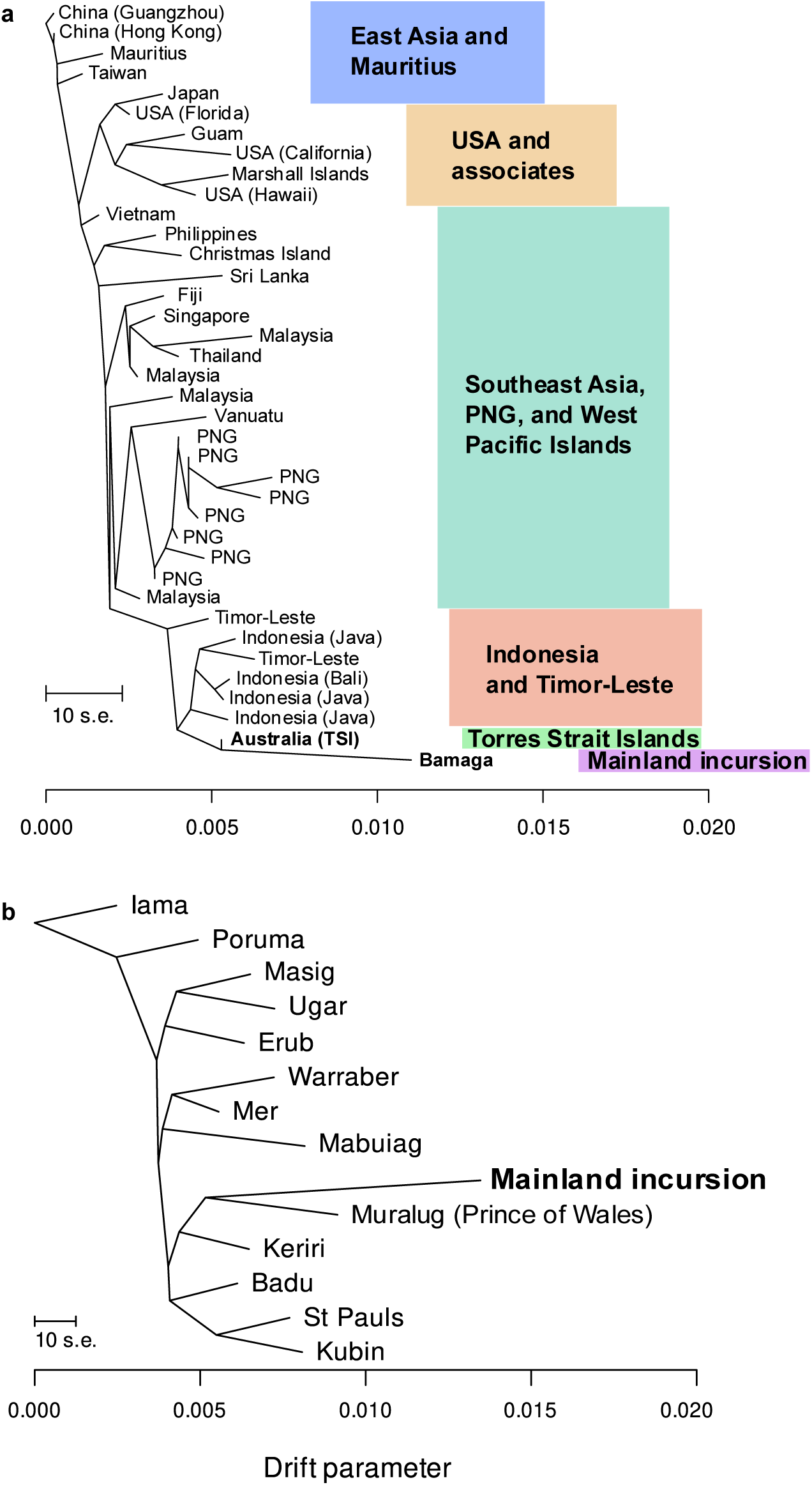
Treemix analysis of mainland incursion origins. (a) Treemix of global *Aedes albopictus* population samples. Treemix was run 100 times, analysing blocks of 100 SNPs. All 100 runs placed the mainland incursion with the Torres Strait Islands. The tree with the highest log likelihood is shown here. The Torres Strait Islands population sample used here contained individuals from across the islands and rarified to n=24. (b) Treemix of Torres Strait Island population samples. Treemix was run 100 times, analysing blocks of 500 SNPs. All 100 runs placed the mainland incursion with Muralug(Prince of Wales Island). The tree with the highest log likelihood is shown here.

#### Detection lag in mainland incursion

The internal branch leading to the mainland incursion samples was much longer than the internal branches within the other TSI populations, suggesting that the strong drift experienced by the mainland incursion is not that from the original TSI invasion in 2004, but rather reflects a recent bottleneck. If so, this is evidence that the incursion on the Australian mainland occurred several months before detection, and several generations had passed between incursion and detection.

This hypothesis found further support in analysis of Tajima’s D and pairwise relatedness (Fig. 3). For each parameter, we compared the mainland incursion against the Muralug reference population, rarifying the latter to n=13 to remove potential sample size biases. Tajima’s D has an expected value of 0 for populations evolving at equilibrium, but positive Tajima’s D values are expected after a bottleneck, when the site frequency spectrum is biased towards medium and high frequency alleles (Tajima, 1989). Tajima’s D was close to 0 for the Muralug reference population (x̅ = 0.0145, s.d. = 0.899), but positive for the mainland incursion (x̅ = 0. 4643, s.d. = 0.872) (t-Test: t = 14.004, df = 3039.9, p-value < 1e-16) (Fig. 3a).

**Figure 3.**
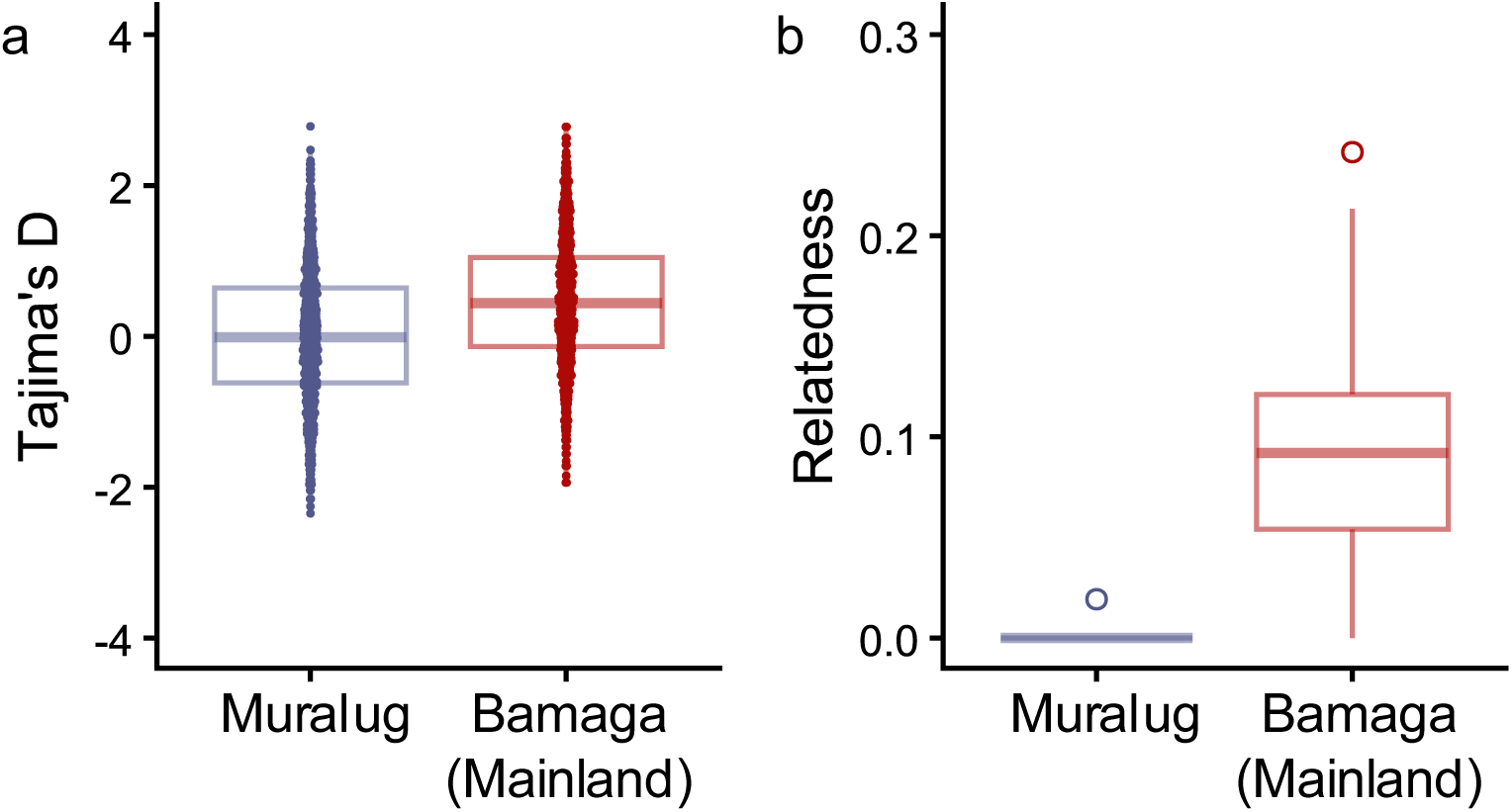
Population genetic comparisons of old (Muralug) and new (Bamaga) invasions. Both populations are rarified to n=13 individuals. (a) Tajima’s D, calculated in 100 kb windows. Muralug, x̅ = 0.0145; Bamagxa, = 0.4643. t-Test: t = -14.004, df = 3039.9, p-value < 2.2e-16. (b) Pairwise relatedness.

Pairwise relatedness is expected to be higher following an invasion bottleneck, as all individuals are derived from the same genetic propagule, resulting to an increase in population-level inbreeding. In line with these predictions, we observed much higher mean relatedness on the mainland (x̅ = 0. 0895, s.d. = 0.0533) than on Muralug (x̅ = 0. 0002, s.d. = 0.0022) (Fig. 3b). The higher mean relatedness was not due to a few highly inbred pairs, as the highest relatedness was 0.242, which is at the level of a 1^st^ order relationship (e.g. parent-offspring or full-sib) in an outbred population. Rather, it represents a higher level of relatedness among most individuals in the sample.

### Island incursions

#### Origins of island incursions

Unlike the mainland incursion, PCAs of the island incursions did not indicate strong drift patterns with samples plotting far apart from others (Fig. S1b,c; c.f. S1a). Instead, most incursion samples clustered with reference samples from the inner islands Keriri and Muralug (Fig. S1b,c). We inferred origins using Locator, where the cloud of points from the 1000 bootstrap replicates of each incursive sample either clustered around one specific island, or around either of the two inner islands (Keriri or Muralug, Fig. 1), or was of an unknown origin. Figure 4 shows examples of these. Results are summarized in Table 1. Detailed results are in Figs S2—S64.

**Figure 4.**
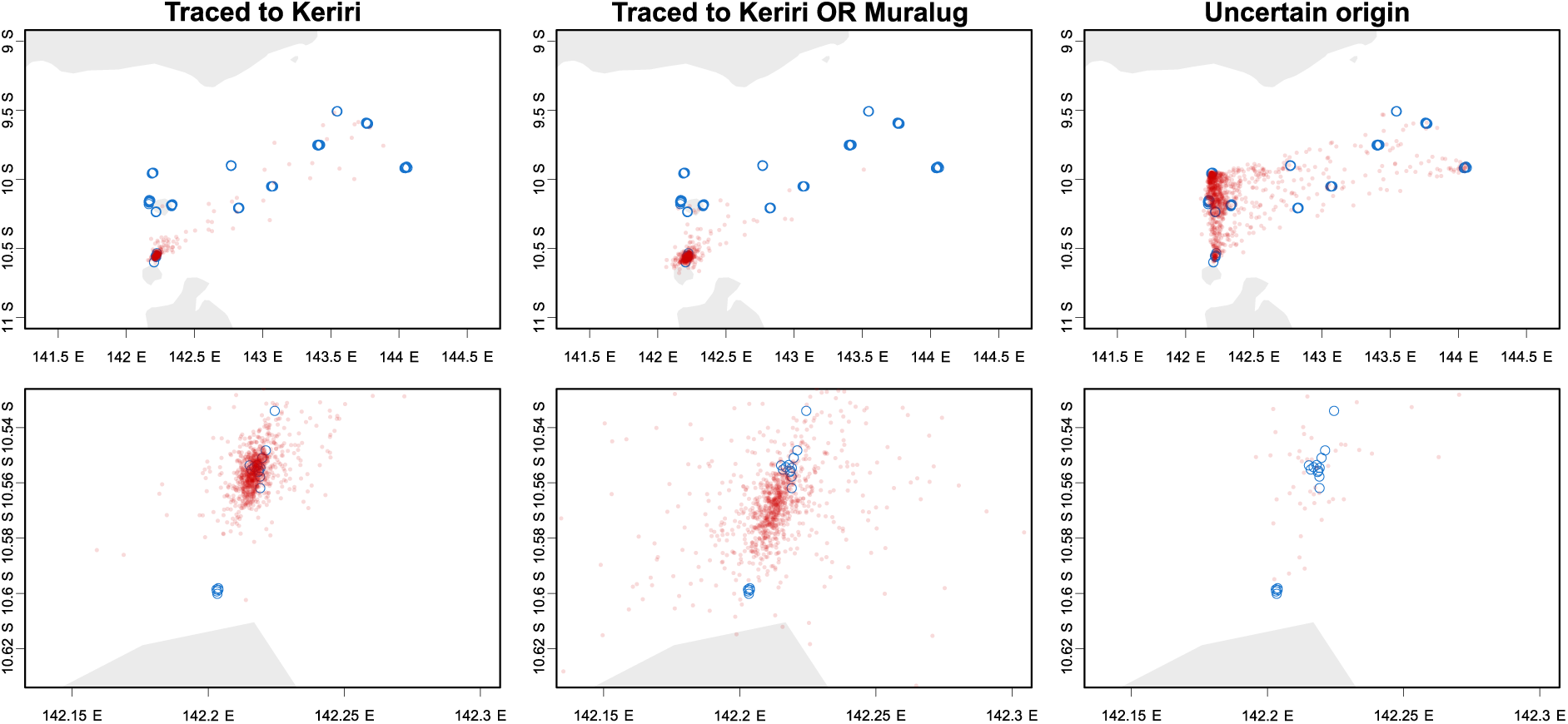
Example categorisation of Locator origin tracing results. Columns show tracing results for three incursions, traced confidently to Keriri (left), to either Keriri or Muralug (centre), or of uncertain origin (right). Blue circles show reference samples locations. Red circles show n=1000 bootstrap replicates of inferred origin. Top plot shows all Torres Strait Islands, bottom plot is the same analysis showing details around Keriri (top blue cluster) and Muralug (bottom blue cluster). Results for all incursions are in Figs S2—S64.

**Table 1.** Summary of tracing results for incursions detected on Ngurapai (Horn) and Waiben (Thursday) islands. Incursions from 2019-2023 were sequenced with ddRAD sequencing, while those from 2025-26 used whole genome sequencing. Keriri OR Muralug refers to incursions with a clear origin within the inner islands but not specific to either island. Detailed tracing results are in Figs S2—S64.

| Year | Island of detection | Keriri (Hammond) | Muralug (Prince of Wales) | Keriri OR Muralug | Moa (Banks) | Badu (Mulgrave) | Uncertain |
|---|---|---|---|---|---|---|---|
| 2019 | Ngurapai | 3 |  |  |  |  | 1 |
| 2021 | Ngurapai |  | 2 |  |  |  | 1 |
|  | Waiben |  | 2 |  | 1 |  |  |
| 2022 | Ngurapai | 2 | 2 | 1 |  |  |  |
|  | Waiben | 1 | 2 | 1 |  |  |  |
| 2023 | Ngurapai |  |  |  |  | 1 |  |
|  | Waiben | 10 | 6 | 7 | 1 |  | 3 |
| 2025 | Waiben | 3 | 3 | 1 |  |  |  |
| 2026 | Ngurapai | 2 | 5 |  |  |  | 1 |
|  | Waiben | 1 |  |  |  |  |  |

Several patterns emerge from these results. Firstly, the vast majority of the n=63 incursions were confidently traced to the inner islands Keriri and Muralug (n=54). Only n=3 were specifically traced to outer islands. These patterns were consistent across years, and Ngurapai and Waiben both received similar proportions of incursions from each of Keriri and Muralug. All 63 incursions were traced to the TSI.

Previous analyses in this system had much lower success rates for tracing incursions. Incursions collected in 2019 had 50% successful tracing (Schmidt et al., 2021), while those from 2021–2022 had 53.3% success (Schmidt et al., 2023). Here, our reanalysis was bolstered by the inclusion of the Muralug reference samples and the current contiguous reference assembly for *Ae. albopictus*. This led to increases to 75% successful tracing in 2019 and 80% for 2021–2022; this latter statistic increases to 93.3% if successful tracing included “Keriri OR Muralug”. Overall, considering both new and reanalysed incursions, 74.6% of incursions were successfully traced, which increased to 90.5% if successful tracing included “Keriri OR Muralug”.

#### Composite analysis of whole genome and reduced representation sequence data

The incursion results also allowed us to compare tracing success when incursions were sequenced with ddRADseq vs with WGS. As the TSI reference sequences were all produced with ddRADseq, no benefit would be expected from the additional sites available through WGS, as these were not genotyped in the reference databank. Therefore, this comparison was effectively assessing whether using WGS for incursions would negatively affect tracing. We observed no decrease in tracing using WGS. For ddRADseq, 70.2% of incursions were successfully traced, which increased to 89.4% if successful tracing included “Keriri OR Muralug”. For WGS, 87.5% of incursions were successfully traced, which increased to 93.8% if successful tracing included “Keriri OR Muralug”.

Although the above results suggest that the transition to WGS for incursion samples did not negatively affect tracing to the ddRAD databank, there was some loss of genetic structure observed in PCAs (Fig. S1b, c.f. S1c). This was likely caused by the composite WGS C ddRAD genotyping and filtering resulting in fewer retained SNPs (n=14,812) compared with ddRAD only (n=36,081). When only ddRAD samples were analysed, processing could be through the Stacks pipeline which is optimized specifically for RADseq (Rochette et al., 2019). Processing both data types together in GATK (Van der Auwera C O’Connor, 2020) likely resulted in many SNPs failing to pass filters. Although we could have reprocessed the composite dataset using minimal filtering (e.g. without GATK hard filters) to retain more SNPs, this could introduce new, unknown biases. As tracing results did not seem adversely affected by the reduction in SNPs, we kept the standard GATK filters out of caution.

#### Independence of incursions

When incursive individuals are genetically related, this suggests they are likely not from independent incursions. Instead, they may have dispersed together on the same boat (as eggs, larvae, or adults), or share a parent that dispersed in a previous generation. We identified four kinship groups in the 63 incursion samples from 2019–2026. Within each kinship group, all individuals were collected at the same time and either in the exact same location (three groups) or at properties next door to one another (one group).

For each kinship group, origin tracing produced no discordant results. Either all individuals were assigned the same origin (two groups), or contained a single individual of uncertain assignment (two groups). We posited that these uncertain assignments could be assigned specific origins by assigning them the same origin as their kin. Using these imputed origins, this increased the proportion of successfully traced incursions to 77.8%, which increased to 92.1% if successful tracing included “Keriri OR Muralug”.

## Discussion

Proactive management of invasive species aims to prevent incursions and to detect and eradicate incursions before establishment. Compared with reactive management of established invasions, proactive management can be cheaper, reduces local damages in the short term, and buys time to prepare for later establishment, but is nevertheless used much less frequently (Ahmed et al., 2022; Cuthbert et al., 2022). The TSI biosecurity zone in northeastern Australia represents a highly successful example of proactive management. Since 2004, when *Ae. albopictus* first invaded the TSI, proactive biosecurity has prevented the species spreading to establish on the Australian mainland approximately 20 km to the south. Over this same period of time, *Ae. albopictus* has spread rapidly through Europe where it has reached a current northwards expansion rate of 60.3 km per year (Farooq et al., 2025). The 2026 incursion in Bamaga marked the first discovery of a thriving *Ae. albopictus* population on the Australian mainland, where it had previously only been detected as isolated interceptions at port areas associated with international cargo. This study has described the detection and evolutionary analysis of this population prior to effective vector control intervention. Using other, isolated incursions in the TSI for comparison, we infer that this population had likely been present on the mainland for at least two generations. Given that *Ae. albopictus* spread very quickly through the TSI following its initial establishment (Ritchie et al., 2006), this detection in Bamaga may have come just quickly enough to stop it spreading throughout Cape York communities.

The Bamaga incursion, as well as 22–32 of the 63 incursions into the inner TSI, had an origin in Muralug (Prince of Wales Island). These more recent incursions, from 2025-26, thus follow similar patterns to previous years, where most dispersal into the inner islands is from other inner islands (Keriri and Muralug) with occasional dispersal from outer islands. Critically for this study, Muralug samples were collected and added to the databank in 2023. This led to a major improvement in tracing outcomes, and demonstrates the critical importance of capturing as full a range of genetic variation possible when performing population assignment and other population genetic tests. Reanalysis of previously published tracing results of n=20 incursions (Schmidt et al., 2021, 2023) identified Muralug as the origin of 9 incursions previously characterized as from Keriri (n=3) or an unknown location (n=6). Following the inclusion of Muralug in the databank, tracing success improved by >50%.

While missing populations had clear negative effects on tracing, we found no negative effects from transitioning from ddRAD sequencing of incursions to whole genome sequencing. Although structure was weaker in PCAs (Fig. S1b,c), tracing success remained equivalent across data types (Table 1). This is a promising outcome for combining datasets produced using different sequencing types, including other work on *Ae. albopictus* using SNP chips (Cosme et al., 2024) that could also be analysed alongside WGS databanks (Crawford et al., 2023; Seok et al., 2025). However, such composite datasets require consideration of ascertainment bias, where using only SNPs called in one of the two datasets will miss much genetic variation in the other dataset. Here this was not an issue, as this second dataset contained only the incursion samples, which were of the same gene pool as the dataset in which SNPs were called (the reference dataset).

The origin tracing work in this study builds on work conducted over seven years (2019-2026), with specific aims to inform the management of *Ae. albopictus* in the TSI and the broader Northern Peninsula Area. The results here are informative to these aims as follows. The absence of inbreeding among the specimens found on Waiben and Muralug over several years confirms that the vector control strategies in use have been effective at repeatedly eradicating *Ae. albopictus* and preventing its establishment on the two islands, despite frequent reinvasion from nearby islands. With genomic evidence demonstrating that most incursions on Waiben and Ngurapai originate from Muralug and Keriri islands, and with the Bamaga incursion also linked to Muralug, the feasibility of expanding the vector control program to suppress *Ae. albopictus* populations on Muralug and Keriri is under consideration. This potential expansion, which would effectively eradicate *Ae. albopictus* from the inner TSI (Fig. 1a), could help reduce the incursion pressure coming from invaded islands. Considering that the Bamaga *Ae. albopictus* incursion was detected after several generations of breeding, the program would benefit from increasing the frequency of monitoring throughout the Northern Peninsula Area communities, to facilitate early detection. Early detection is essential in Northern Peninsula Area, because *Ae. albopictus* can spread quickly, and eradication may not be possible once the species is firmly established in the region.

The high tracing precision and success, maintained over eight years, shows the TSI invasion of *Ae. albopictus* is a highly tractable system for incursion tracing. One advantage the TSI possesses is that the islands enforce discretization on the system, so that islands develop discrete structure from one another, and the distance between islands maintains this structure even though dispersal is fairly common between islands (Schmidt et al., 2021). Origin tracing in the TSI has also benefited from a sufficient genetic variation in *Ae. albopictus* and sufficient drift during each island establishment, which together have allowed each island to develop its own distinct genetic signature; this pattern was observed even shortly after invasion (Maynard et al., 2017) and remained consistent over the following decade (Schmidt et al., 2021). However, tracing might be less precise in pest systems with more dispersal between locations, such as seen in some *Aedes* (Paris et al., 2023; Schmidt et al., 2025; Webb C Russell, 2019) and *Anopheles* species (Saddler et al., 2019), or where continuity of habitat leads to more continuous genetic structure. In such cases, genomic databanks may be underpowered, or may lose their power more rapidly, as the present-day populations cease to resemble the populations sequenced in the databanks. For systems where genetic structure is insufficient to distinguish pest populations for tracing, it could be possible for similar approaches to be developed that make use of viruses such as insect-specific viruses (Everts et al., 2026; Hollingsworth et al., 2023). These approaches would exploit the higher evolutionary rate of viruses relative to eukaryotes and bacteria (Everts et al., 2026). Although such approaches would pose additional complications for analysis of RNA viruses, as DNA is much easier than RNA to preserve in what are often challenging biosecurity sampling situations, virus studies may be a way forward for systems where important variation occurs at spatially or temporally fine scales.

## Conclusions

*Aedes albopictus* management in the TSI is an example of successful, ongoing, proactive management of an invasive species. Genomic data have been a vital component of this approach. This paper has shown how ongoing genomic monitoring has been able to identify incursion risks associated with specific islands, and to determine if incursions are being detected sufficiently quickly after dispersal to uninvaded areas. Together these are used by managers to decide if current vector control strategies are working and if the geographical range of control should be expanded. By using a long term focus, including reanalysis of published data, this paper also shows how long term datasets can be useful for confirming consistency of patterns and for improving insights as new material is incorporated.

## Data availability

All sequence data will be made publicly available on NCBI SRA by time of publication.

## Acknowledgements

We acknowledge and greatly appreciate the support and collaboration from the leaders and community of Northern Peninsula Area Regional Council for the success of the vector control program.

TLS was funded by an ARC DECRA (DE230100257). The vector surveys and control in the Northern Peninsula Area and in the Torres Strait were funded by the Australian Government and the Queensland Government through the Cairns Public Health Unit, Torres C Cape Public Health Unit, and Northern Peninsula Area Regional Council.

