## Supplementary material for "Origin tracing and detection lag in the first *Aedes albopictus* invasion of mainland Australia and a long-term genomic dataset of mosquito incursions": Figures S1-S64

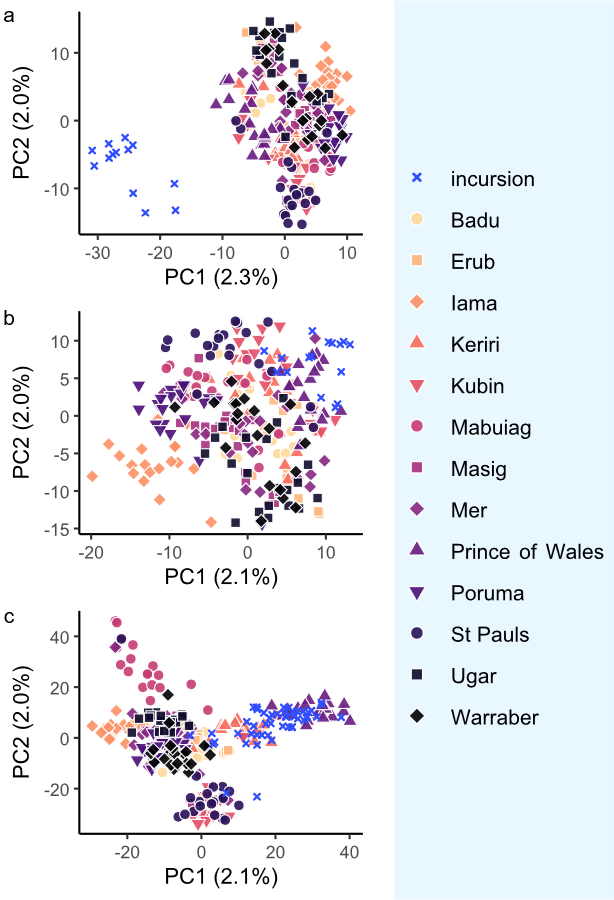

Figure S1. PCAs of TSI reference databank and different sets of incursion samples. (a) Mainland incursion from Bamaga, 2026. (b) Inner TSI incursions 2025-26, using whole genome sequencing. (c) Inner TSI incursions 2018-23, using ddRAD sequencing.

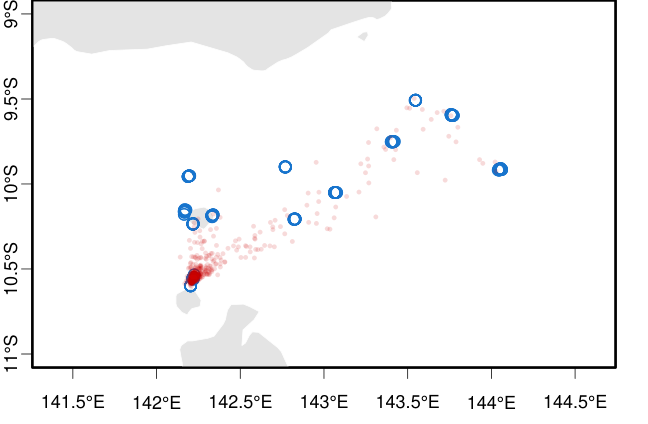

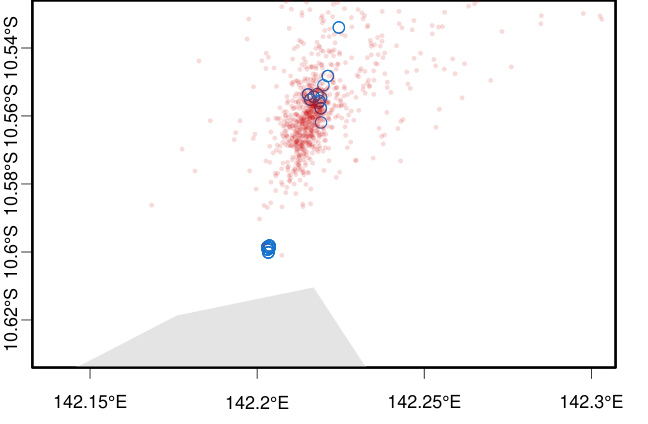

Figure S2. Origin tracing using Locator of sample HI19_001-1, collected on Ngurapai (Horn Island) in 2019. Blue circles show reference samples locations. Red circles show n=1000 bootstrap replicates of inferred origin. Top plot shows all Torres Strait Islands, bottom plot is the same analysis showing details around Keriri (top blue cluster) and Muralug (bottom blue cluster).

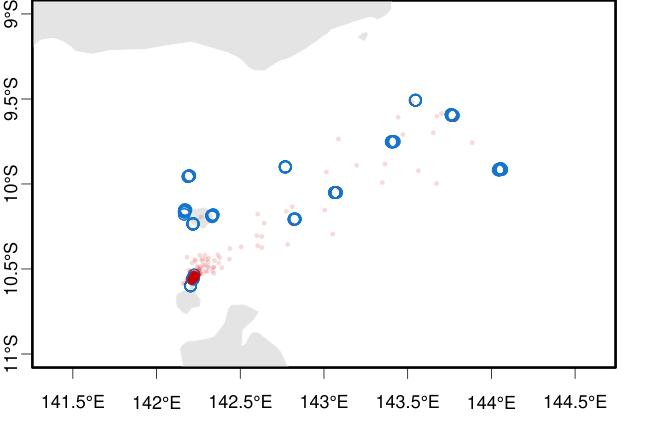

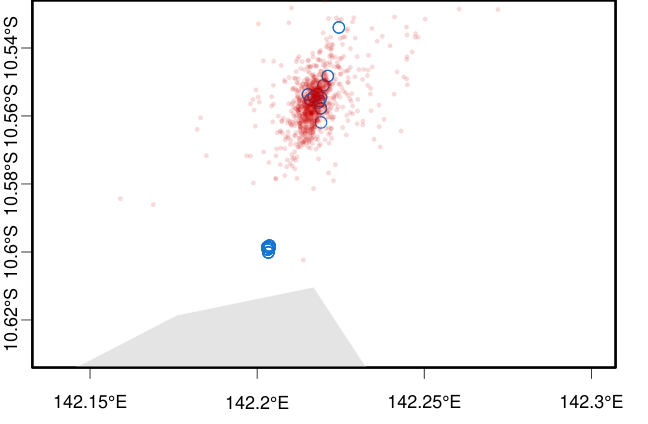

Figure S3. Origin tracing using Locator of sample HI19_001-2, collected on Ngurapai (Horn Island) in 2019. Blue circles show reference samples locations. Red circles show n=1000 bootstrap replicates of inferred origin. Top plot shows all Torres Strait Islands, bottom plot is the same analysis showing details around Keriri (top blue cluster) and Muralug (bottom blue cluster).

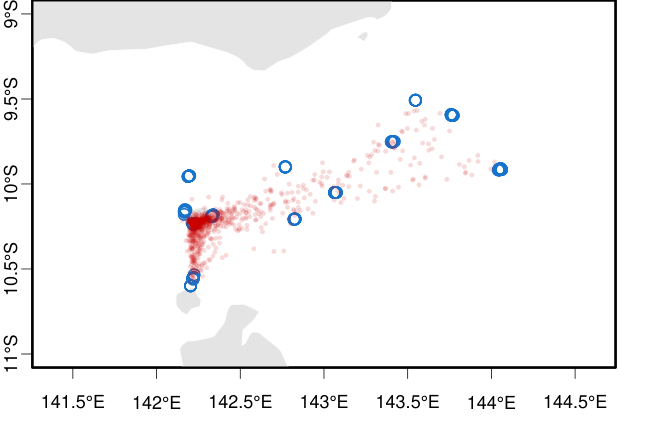

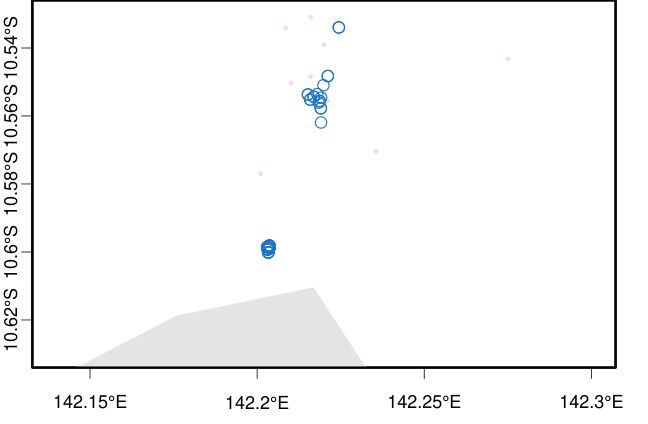

Figure S4. Origin tracing using Locator of sample HI19_058, collected on Ngurapai (Horn Island) in 2019. Blue circles show reference samples locations. Red circles show n=1000 bootstrap replicates of inferred origin. Top plot shows all Torres Strait Islands, bottom plot is the same analysis showing details around Keriri (top blue cluster) and Muralug (bottom blue cluster).

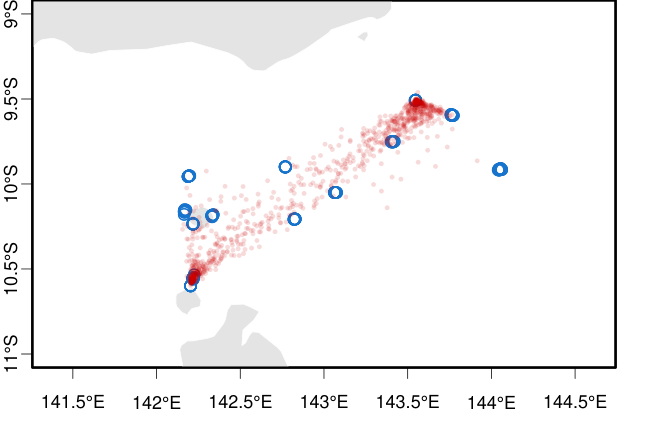

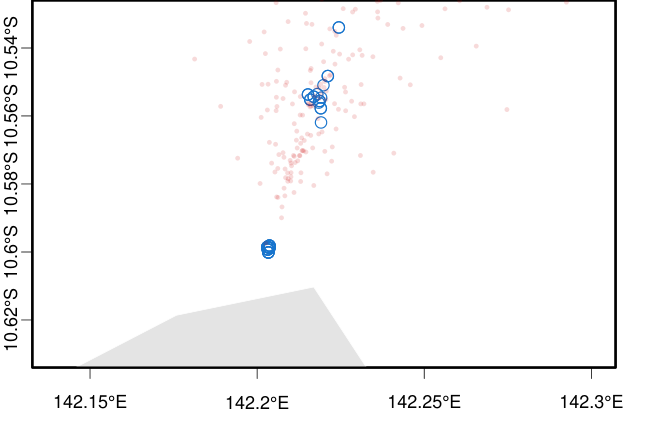

Figure S5. Origin tracing using Locator of sample HI19_095, collected on Ngurapai (Horn Island) in 2019. Blue circles show reference samples locations. Red circles show n=1000 bootstrap replicates of inferred origin. Top plot shows all Torres Strait Islands, bottom plot is the same analysis showing details around Keriri (top blue cluster) and Muralug (bottom blue cluster).

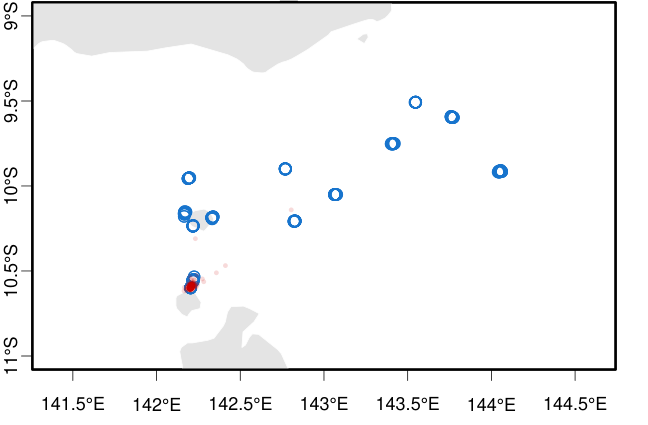

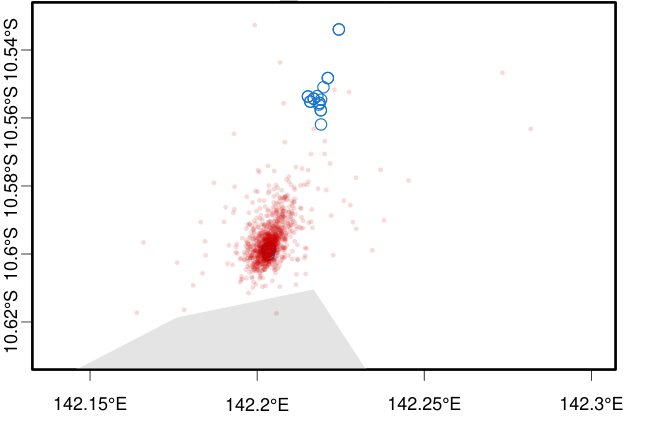

Figure S6. Origin tracing using Locator of sample HI21_001, collected on Ngurapai (Horn Island) in 2021. Blue circles show reference samples locations. Red circles show n=1000 bootstrap replicates of inferred origin. Top plot shows all Torres Strait Islands, bottom plot is the same analysis showing details around Keriri (top blue cluster) and Muralug (bottom blue cluster).

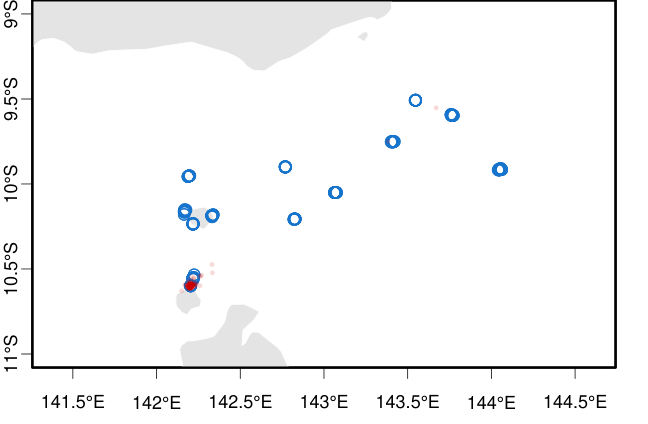

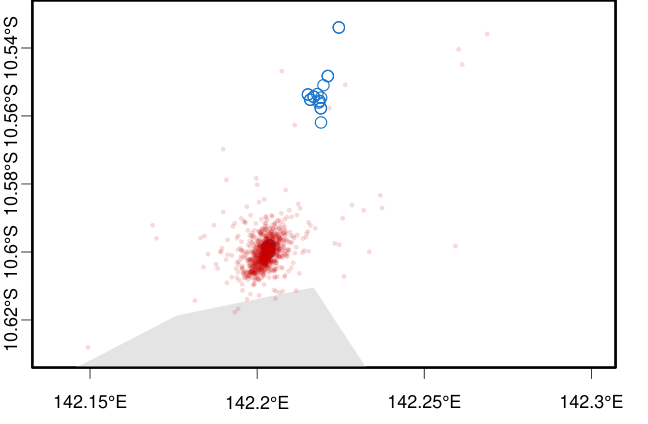

Figure S7. Origin tracing using Locator of sample HI21_009, collected on Ngurapai (Horn Island) in 2021. Blue circles show reference samples locations. Red circles show n=1000 bootstrap replicates of inferred origin. Top plot shows all Torres Strait Islands, bottom plot is the same analysis showing details around Keriri (top blue cluster) and Muralug (bottom blue cluster).

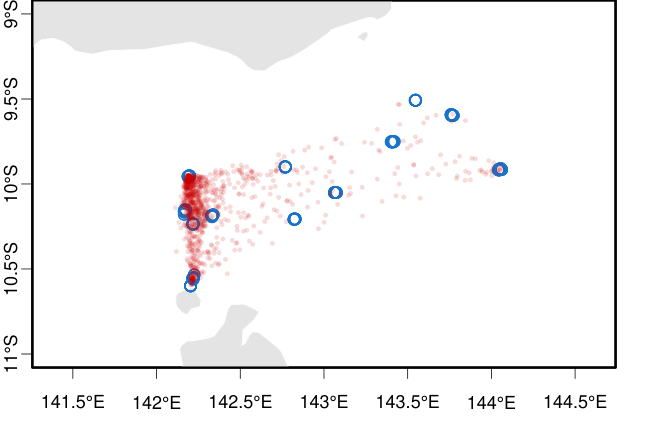

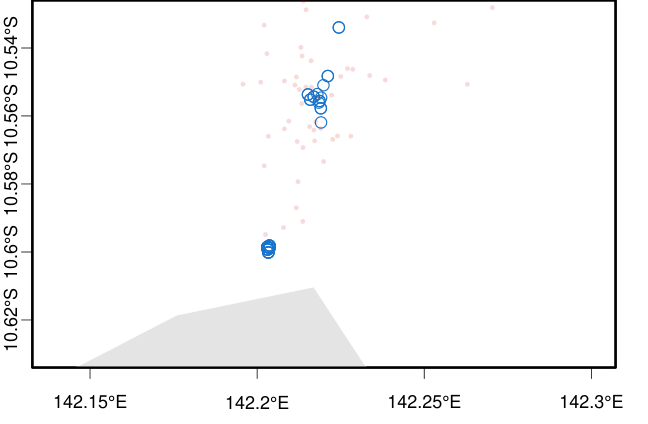

Figure S8. Origin tracing using Locator of sample HI21_041, collected on Ngurapai (Horn Island) in 2021. Blue circles show reference samples locations. Red circles show n=1000 bootstrap replicates of inferred origin. Top plot shows all Torres Strait Islands, bottom plot is the same analysis showing details around Keriri (top blue cluster) and Muralug (bottom blue cluster).

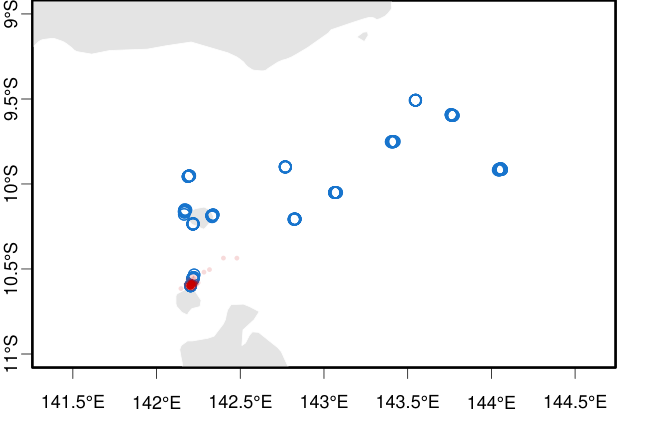

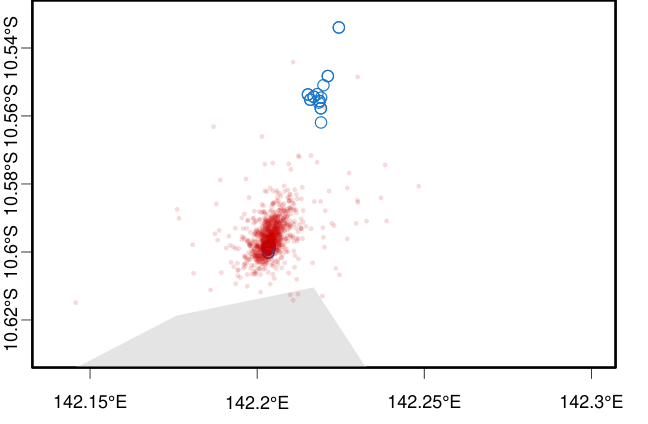

Figure S9. Origin tracing using Locator of sample HI22_030, collected on Ngurapai (Horn Island) in 2022. Blue circles show reference samples locations. Red circles show n=1000 bootstrap replicates of inferred origin. Top plot shows all Torres Strait Islands, bottom plot is the same analysis showing details around Keriri (top blue cluster) and Muralug (bottom blue cluster).

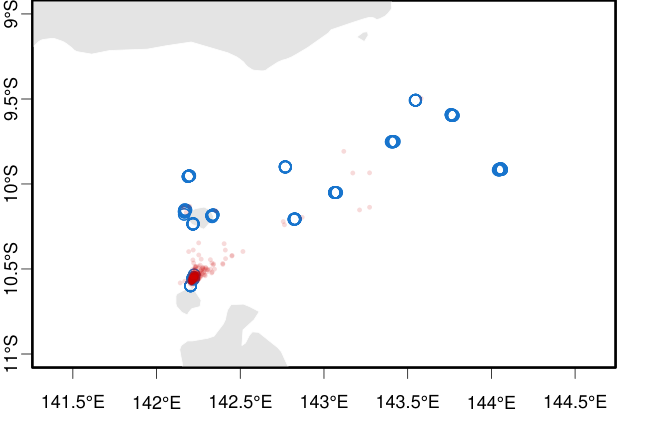

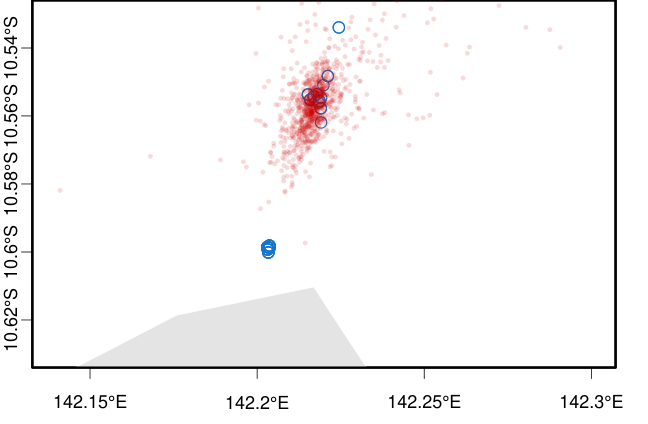

Figure S10. Origin tracing using Locator of sample HI22_055, collected on Ngurapai (Horn Island) in 2022. Blue circles show reference samples locations. Red circles show n=1000 bootstrap replicates of inferred origin. Top plot shows all Torres Strait Islands, bottom plot is the same analysis showing details around Keriri (top blue cluster) and Muralug (bottom blue cluster).

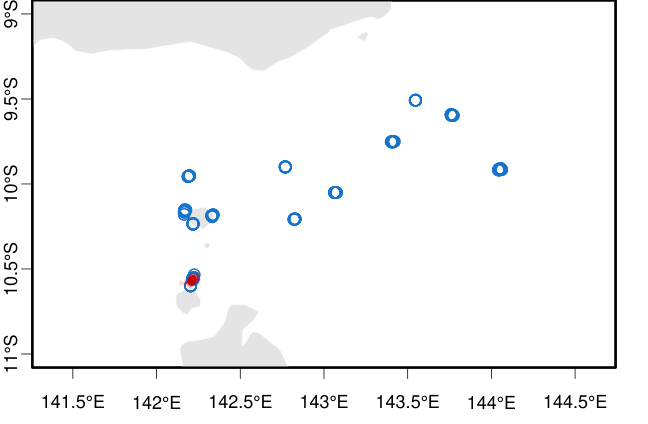

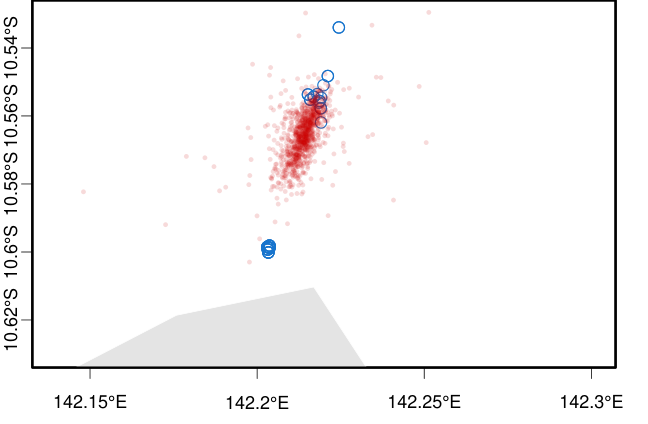

Figure S11. Origin tracing using Locator of sample HI22_065, collected on Ngurapai (Horn Island) in 2022. Blue circles show reference samples locations. Red circles show n=1000 bootstrap replicates of inferred origin. Top plot shows all Torres Strait Islands, bottom plot is the same analysis showing details around Keriri (top blue cluster) and Muralug (bottom blue cluster).

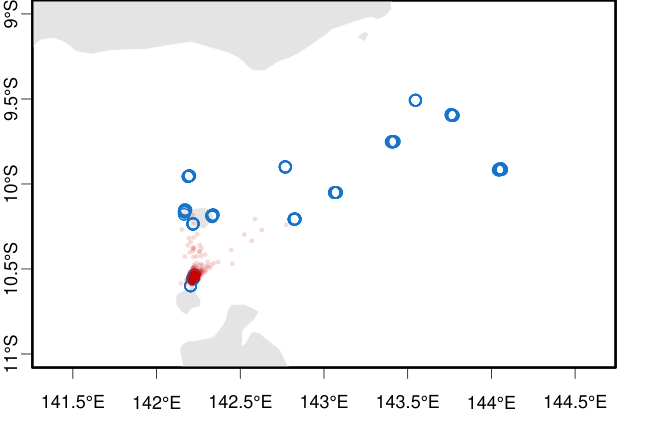

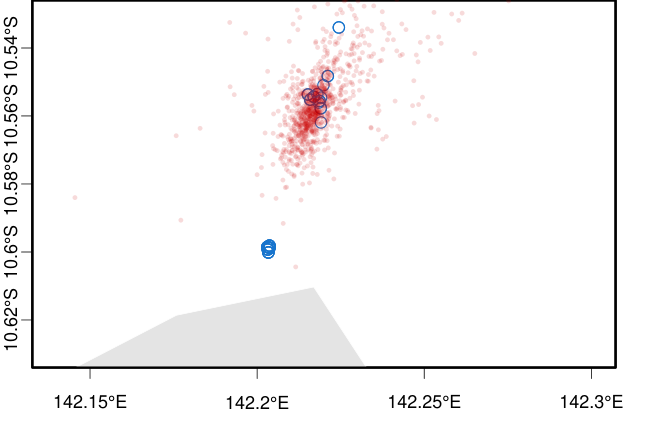

Figure S12. Origin tracing using Locator of sample HI22_077, collected on Ngurapai (Horn Island) in 2022. Blue circles show reference samples locations. Red circles show n=1000 bootstrap replicates of inferred origin. Top plot shows all Torres Strait Islands, bottom plot is the same analysis showing details around Keriri (top blue cluster) and Muralug (bottom blue cluster).

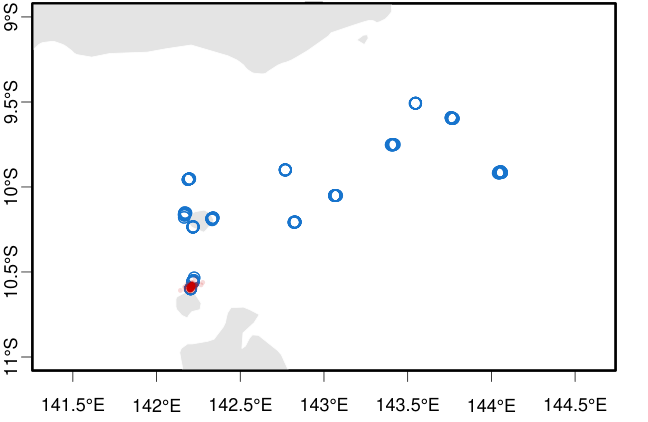

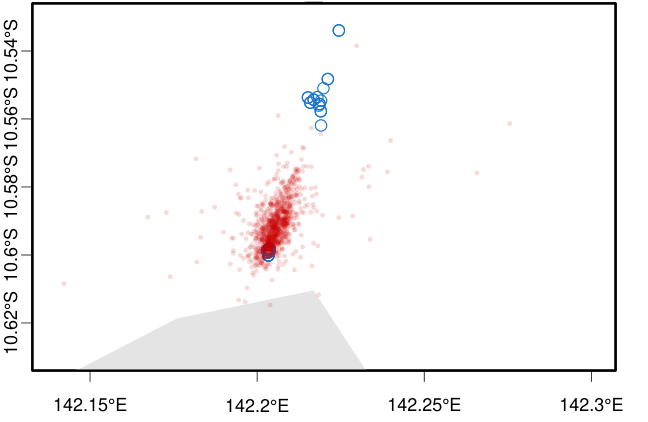

Figure S13. Origin tracing using Locator of sample HI22_097, collected on Ngurapai (Horn Island) in 2022. Blue circles show reference samples locations. Red circles show n=1000 bootstrap replicates of inferred origin. Top plot shows all Torres Strait Islands, bottom plot is the same analysis showing details around Keriri (top blue cluster) and Muralug (bottom blue cluster).

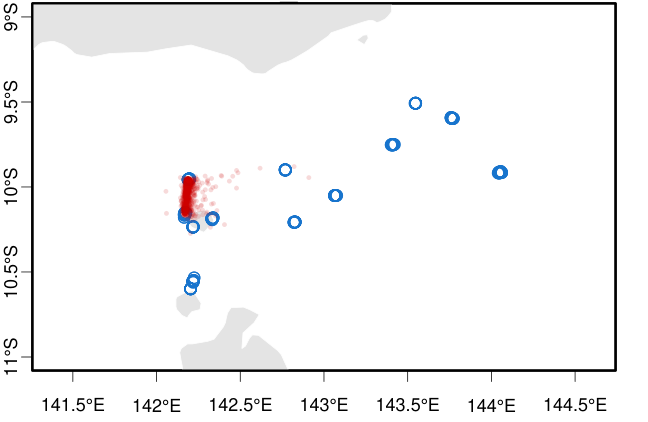

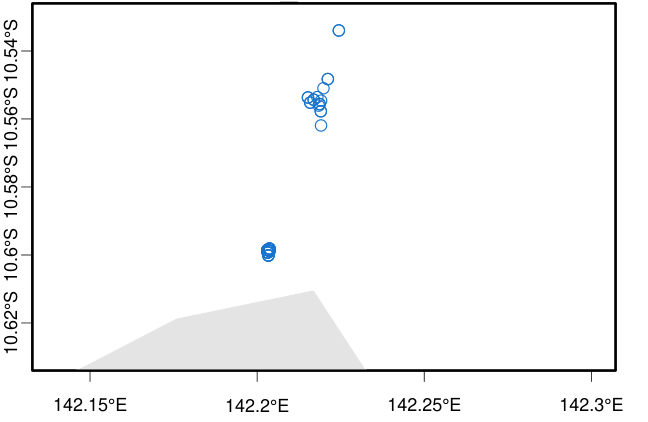

Figure S14. Origin tracing using Locator of sample HI23_006, collected on Ngurapai (Horn Island) in 2023. Blue circles show reference samples locations. Red circles show n=1000 bootstrap replicates of inferred origin. Top plot shows all Torres Strait Islands, bottom plot is the same analysis showing details around Keriri (top blue cluster) and Muralug (bottom blue cluster).

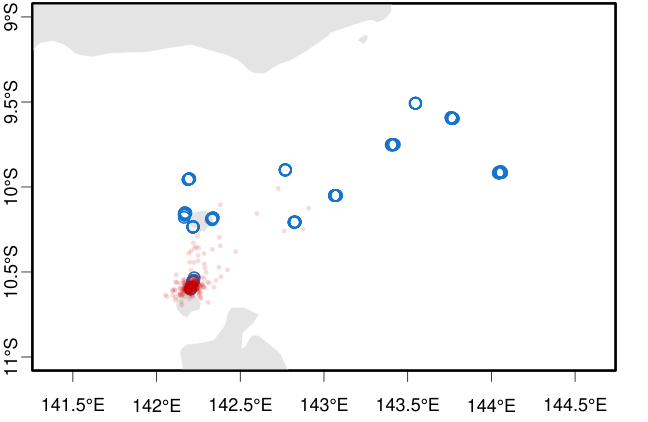

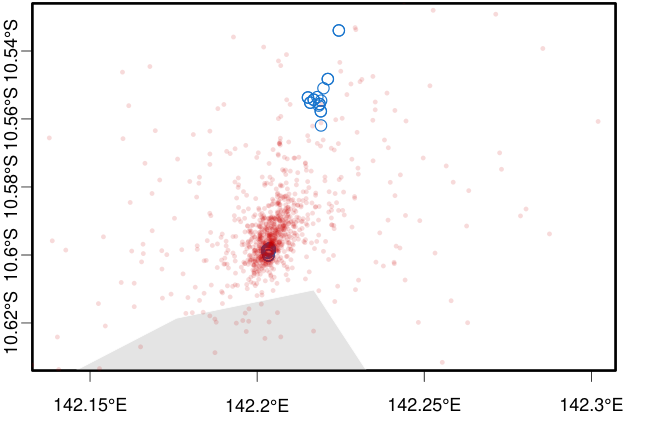

Figure S15. Origin tracing using Locator of sample HI26_001, collected on Ngurapai (Horn Island) in 2026. Blue circles show reference samples locations. Red circles show n=1000 bootstrap replicates of inferred origin. Top plot shows all Torres Strait Islands, bottom plot is the same analysis showing details around Keriri (top blue cluster) and Muralug (bottom blue cluster).

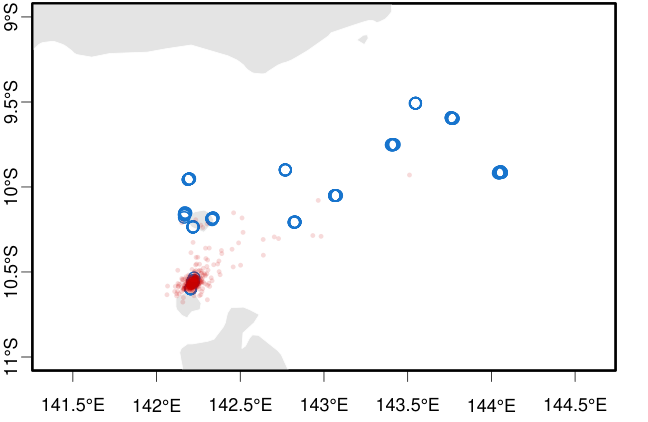

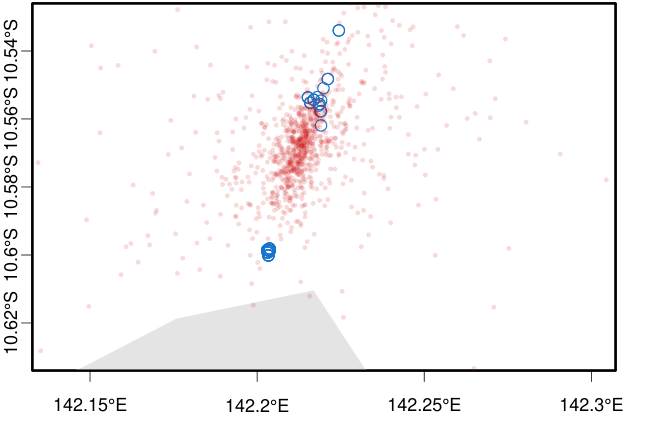

Figure S16. Origin tracing using Locator of sample HI26_110, collected on Ngurapai (Horn Island) in 2026. Blue circles show reference samples locations. Red circles show n=1000 bootstrap replicates of inferred origin. Top plot shows all Torres Strait Islands, bottom plot is the same analysis showing details around Keriri (top blue cluster) and Muralug (bottom blue cluster).

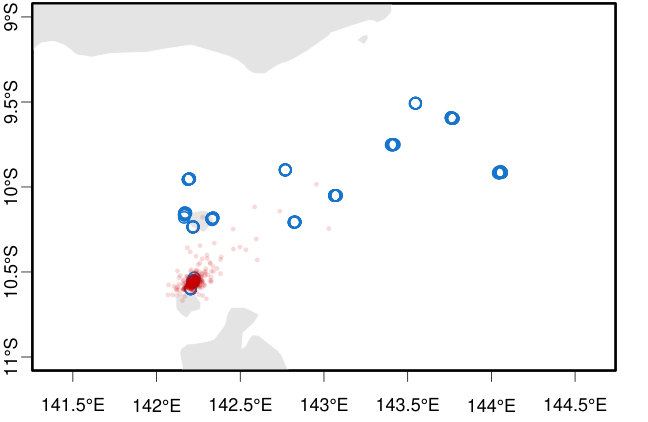

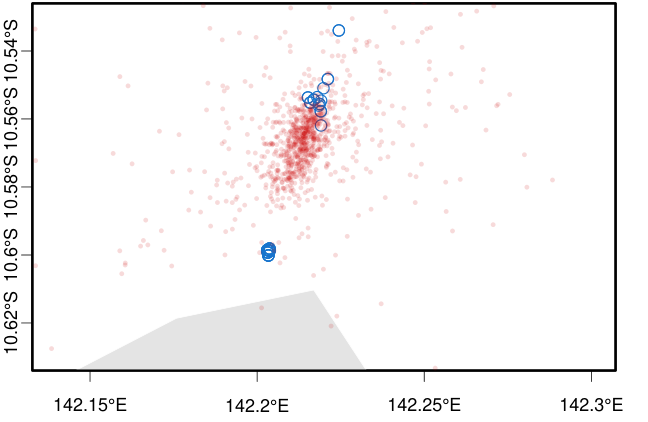

Figure S17. Origin tracing using Locator of sample HI26_034, collected on Ngurapai (Horn Island) in 2026. Blue circles show reference samples locations. Red circles show n=1000 bootstrap replicates of inferred origin. Top plot shows all Torres Strait Islands, bottom plot is the same analysis showing details around Keriri (top blue cluster) and Muralug (bottom blue cluster).

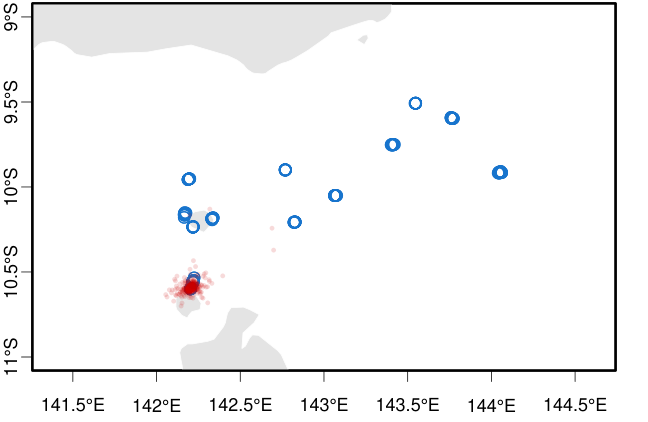

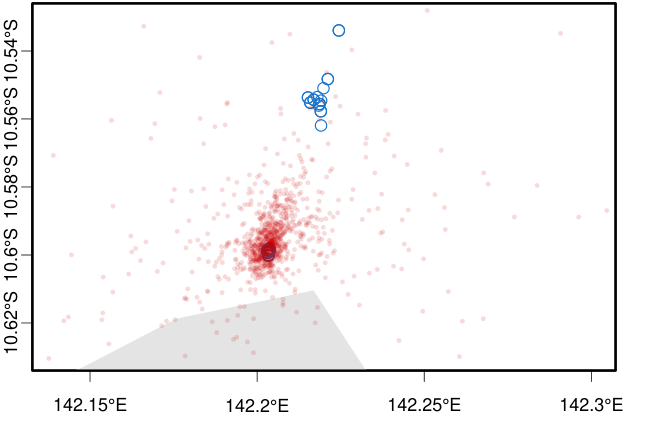

Figure S18. Origin tracing using Locator of sample HI26_065-1, collected on Ngurapai (Horn Island) in 2026. Blue circles show reference samples locations. Red circles show n=1000 bootstrap replicates of inferred origin. Top plot shows all Torres Strait Islands, bottom plot is the same analysis showing details around Keriri (top blue cluster) and Muralug (bottom blue cluster).

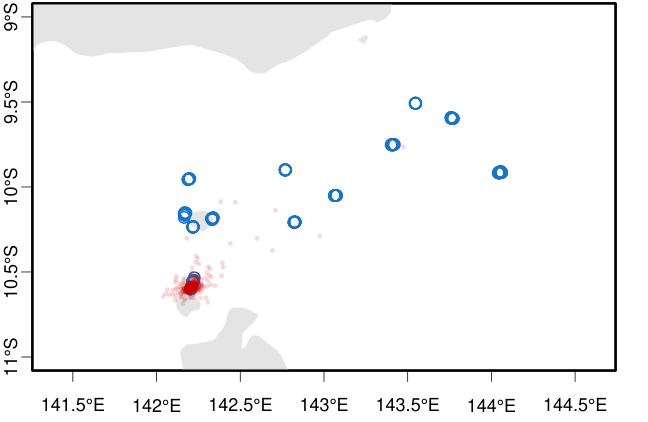

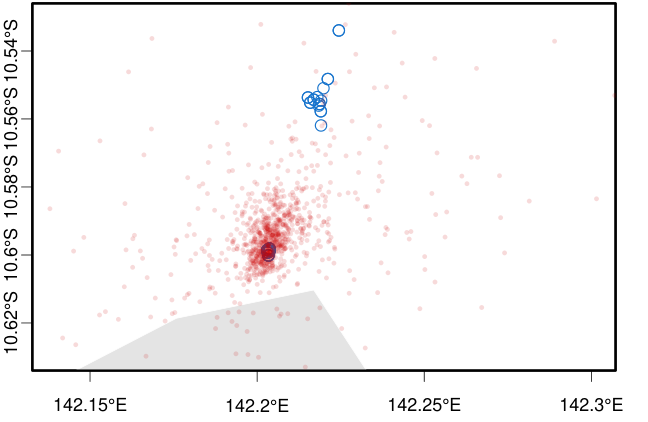

Figure S19. Origin tracing using Locator of sample HI26_065-2, collected on Ngurapai (Horn Island) in 2026. Blue circles show reference samples locations. Red circles show n=1000 bootstrap replicates of inferred origin. Top plot shows all Torres Strait Islands, bottom plot is the same analysis showing details around Keriri (top blue cluster) and Muralug (bottom blue cluster).

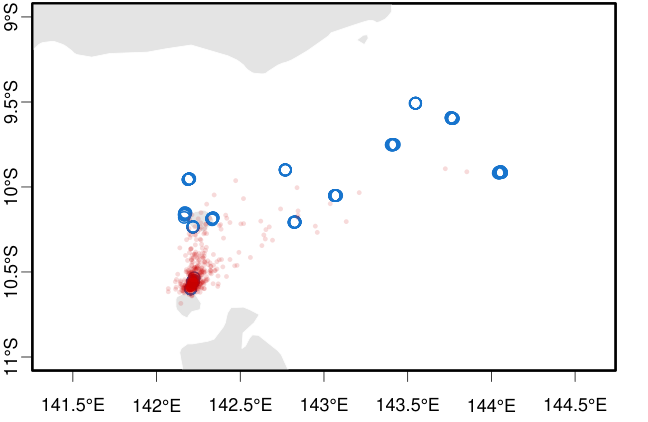

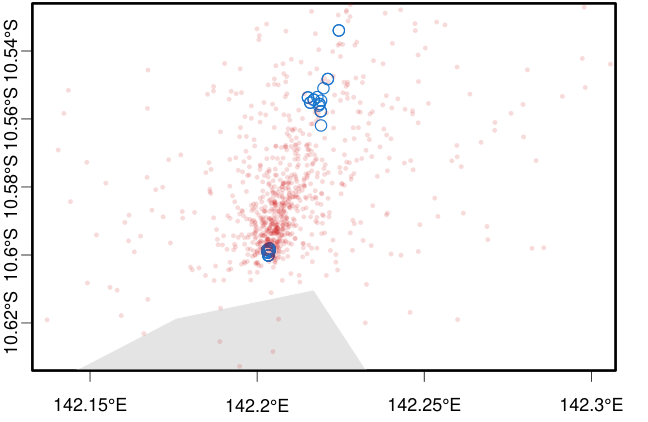

Figure S20. Origin tracing using Locator of sample HI26_074-1, collected on Ngurapai (Horn Island) in 2026. Blue circles show reference samples locations. Red circles show n=1000 bootstrap replicates of inferred origin. Top plot shows all Torres Strait Islands, bottom plot is the same analysis showing details around Keriri (top blue cluster) and Muralug (bottom blue cluster).

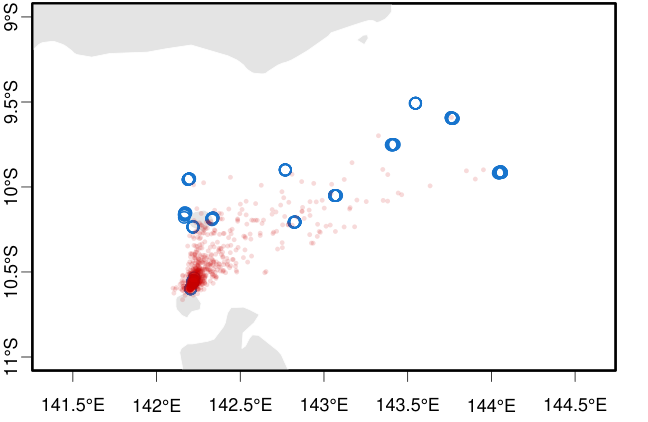

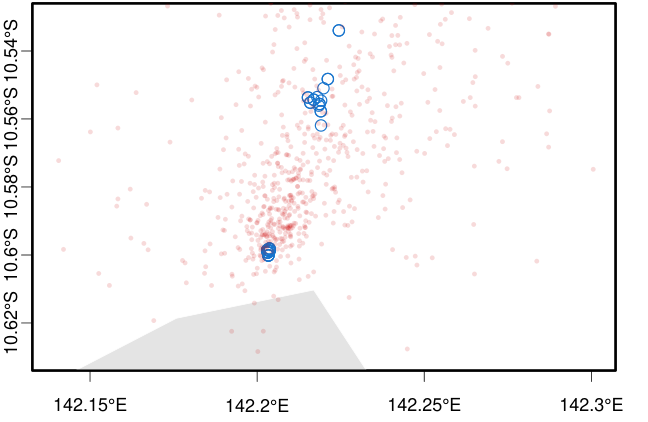

Figure S21. Origin tracing using Locator of sample HI26_074-2, collected on Ngurapai (Horn Island) in 2026. Blue circles show reference samples locations. Red circles show n=1000 bootstrap replicates of inferred origin. Top plot shows all Torres Strait Islands, bottom plot is the same analysis showing details around Keriri (top blue cluster) and Muralug (bottom blue cluster).

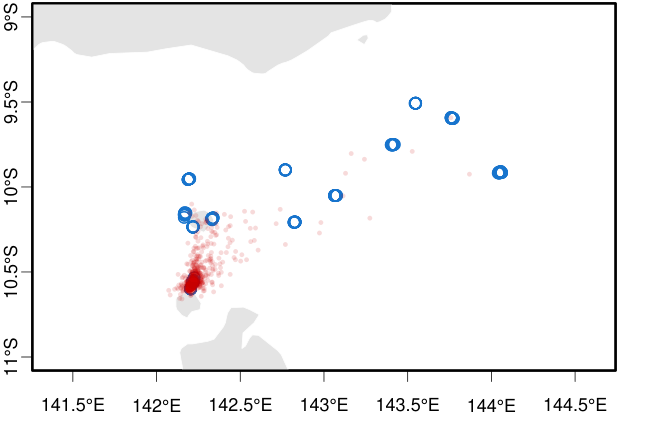

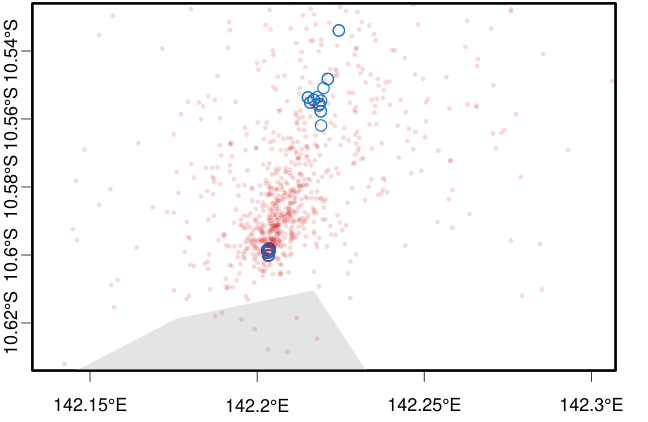

Figure S22. Origin tracing using Locator of sample HI26_074-3, collected on Ngurapai (Horn Island) in 2026. Blue circles show reference samples locations. Red circles show n=1000 bootstrap replicates of inferred origin. Top plot shows all Torres Strait Islands, bottom plot is the same analysis showing details around Keriri (top blue cluster) and Muralug (bottom blue cluster).

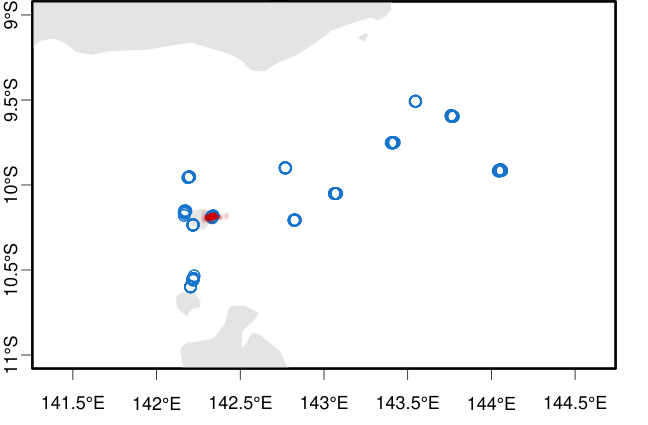

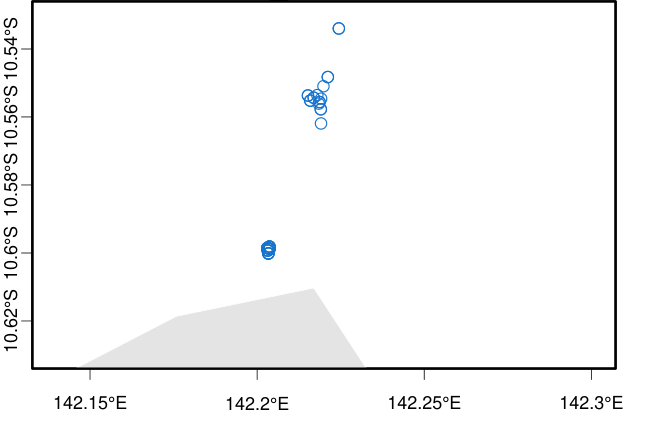

Figure S23. Origin tracing using Locator of sample TI21_030, collected on Waiben (Thursday Island) in 2021. Blue circles show reference samples locations. Red circles show n=1000 bootstrap replicates of inferred origin. Top plot shows all Torres Strait Islands, bottom plot is the same analysis showing details around Keriri (top blue cluster) and Muralug (bottom blue cluster).

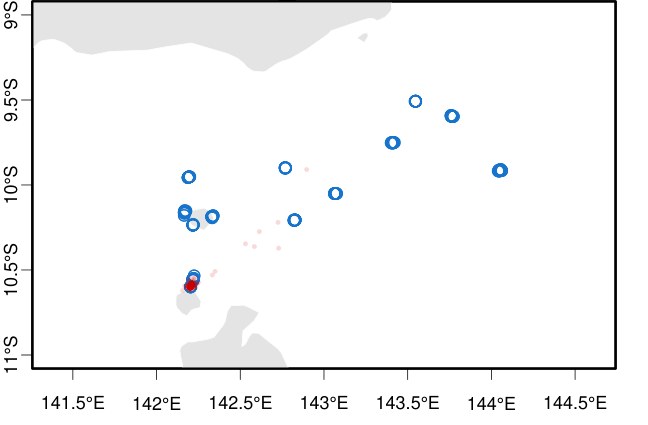

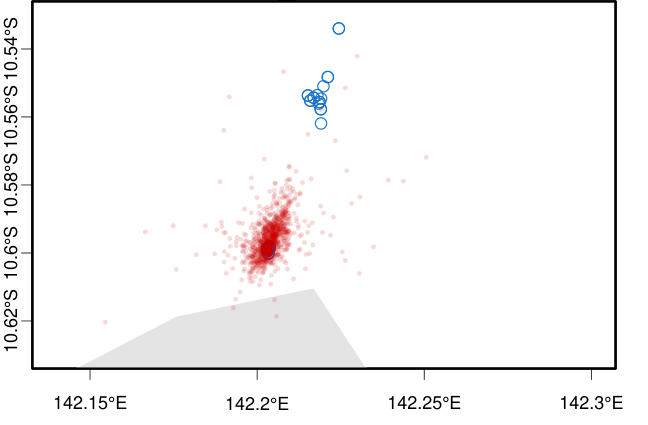

Figure S24. Origin tracing using Locator of sample TI21_041, collected on Waiben (Thursday Island) in 2021. Blue circles show reference samples locations. Red circles show n=1000 bootstrap replicates of inferred origin. Top plot shows all Torres Strait Islands, bottom plot is the same analysis showing details around Keriri (top blue cluster) and Muralug (bottom blue cluster).

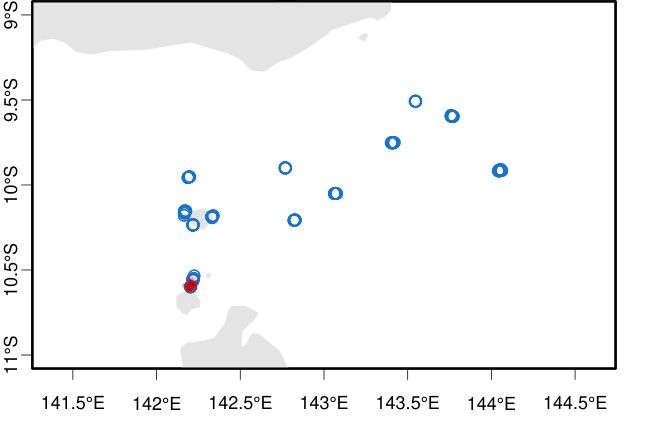

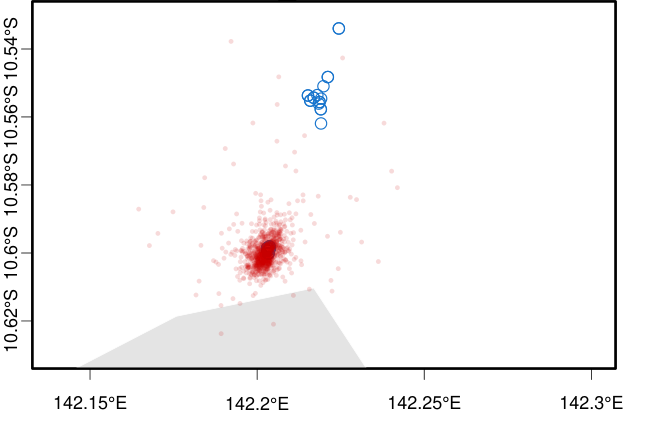

Figure S25. Origin tracing using Locator of sample TI21_139, collected on Waiben (Thursday Island) in 2021. Blue circles show reference samples locations. Red circles show n=1000 bootstrap replicates of inferred origin. Top plot shows all Torres Strait Islands, bottom plot is the same analysis showing details around Keriri (top blue cluster) and Muralug (bottom blue cluster).

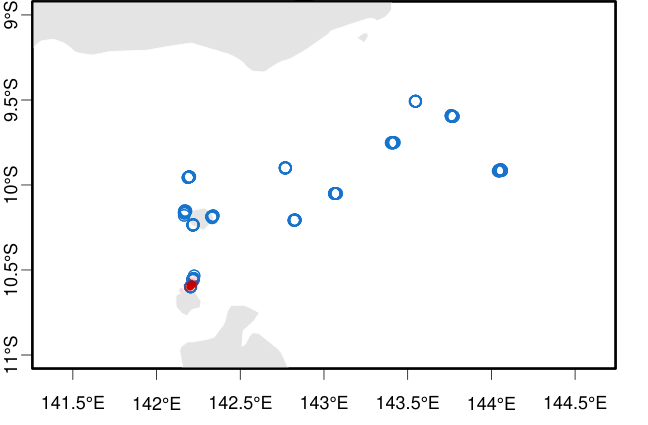

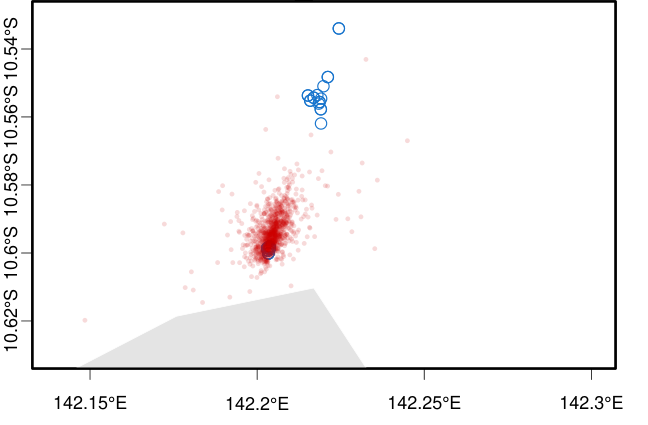

Figure S26. Origin tracing using Locator of sample TI22_152, collected on Waiben (Thursday Island) in 2022. Blue circles show reference samples locations. Red circles show n=1000 bootstrap replicates of inferred origin. Top plot shows all Torres Strait Islands, bottom plot is the same analysis showing details around Keriri (top blue cluster) and Muralug (bottom blue cluster).

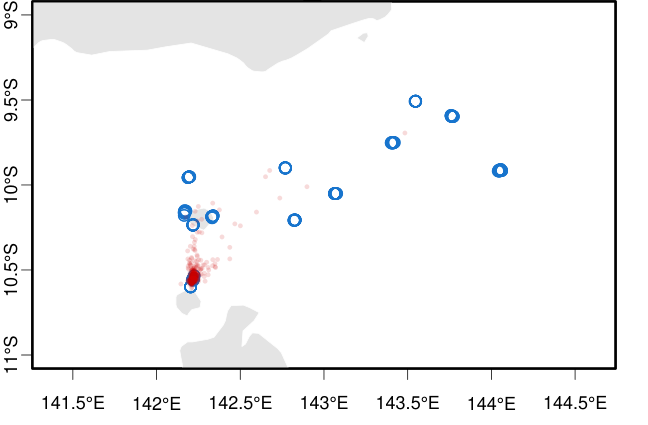

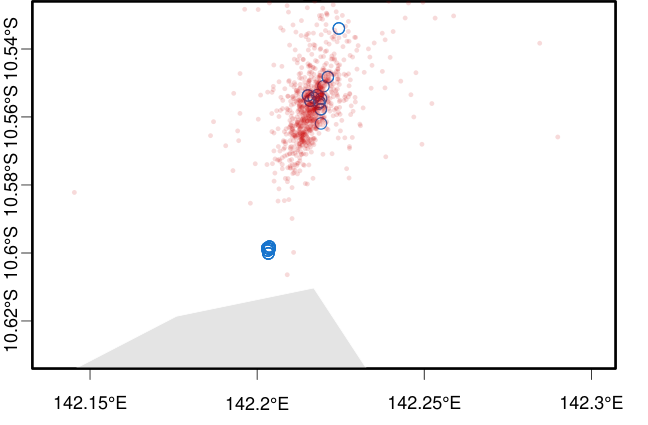

Figure S27. Origin tracing using Locator of sample TI22_016, collected on Waiben (Thursday Island) in 2022. Blue circles show reference samples locations. Red circles show n=1000 bootstrap replicates of inferred origin. Top plot shows all Torres Strait Islands, bottom plot is the same analysis showing details around Keriri (top blue cluster) and Muralug (bottom blue cluster).

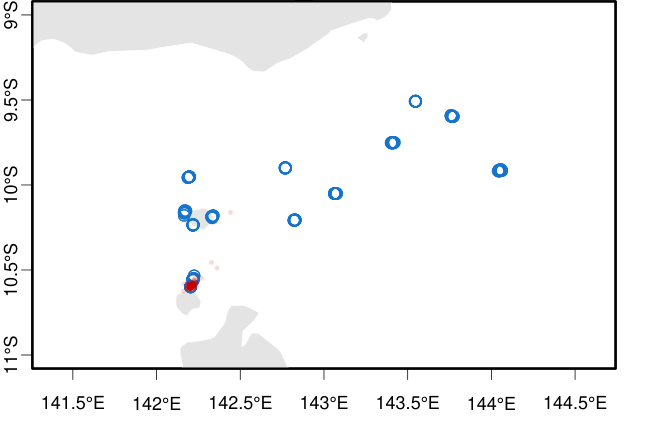

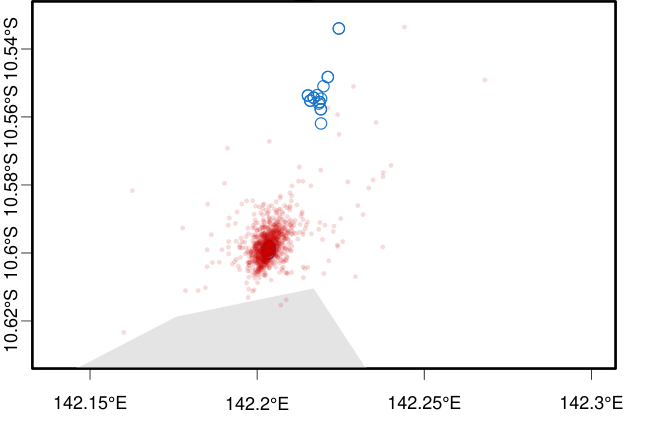

Figure S28. Origin tracing using Locator of sample TI22_170, collected on Waiben (Thursday Island) in 2022. Blue circles show reference samples locations. Red circles show n=1000 bootstrap replicates of inferred origin. Top plot shows all Torres Strait Islands, bottom plot is the same analysis showing details around Keriri (top blue cluster) and Muralug (bottom blue cluster).

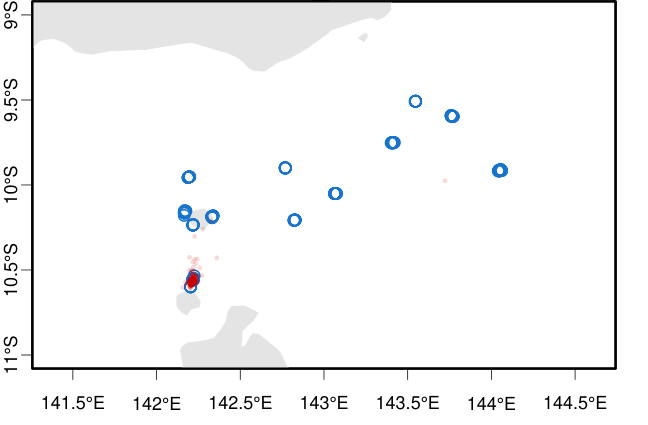

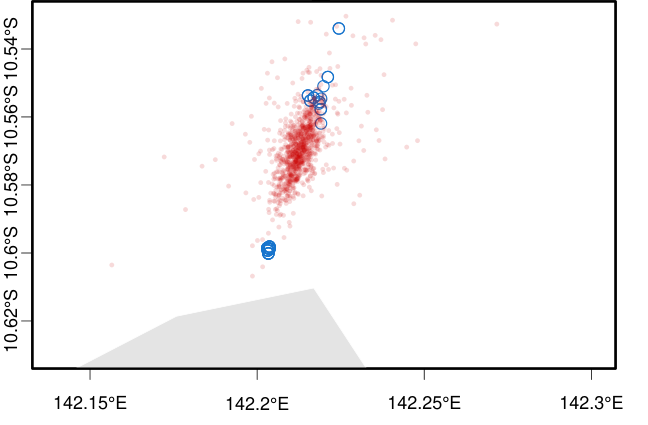

Figure S29. Origin tracing using Locator of sample TI22_009, collected on Waiben (Thursday Island) in 2022. Blue circles show reference samples locations. Red circles show n=1000 bootstrap replicates of inferred origin. Top plot shows all Torres Strait Islands, bottom plot is the same analysis showing details around Keriri (top blue cluster) and Muralug (bottom blue cluster).

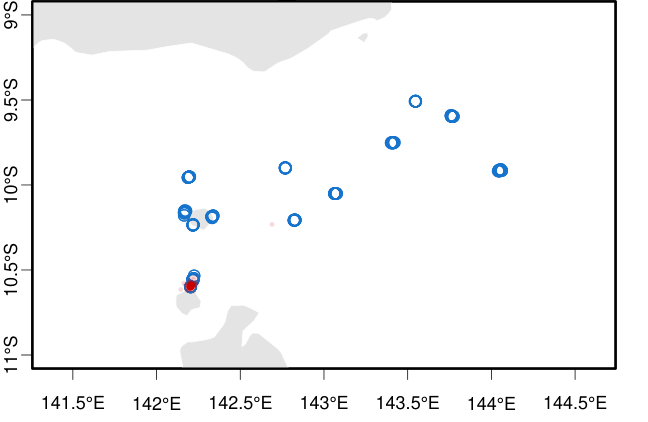

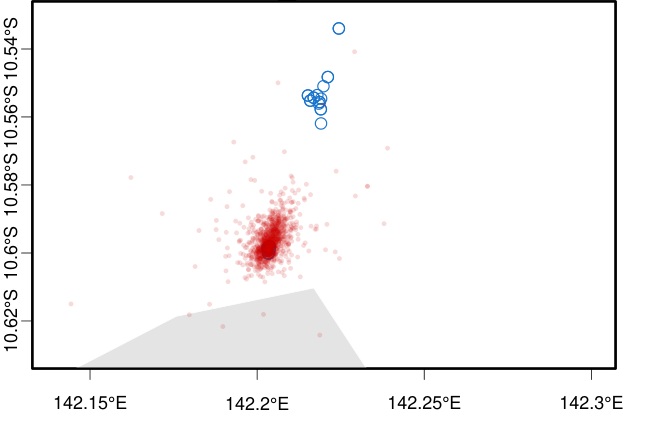

Figure S30. Origin tracing using Locator of sample TI23_043, collected on Waiben (Thursday Island) in 2023. Blue circles show reference samples locations. Red circles show n=1000 bootstrap replicates of inferred origin. Top plot shows all Torres Strait Islands, bottom plot is the same analysis showing details around Keriri (top blue cluster) and Muralug (bottom blue cluster).

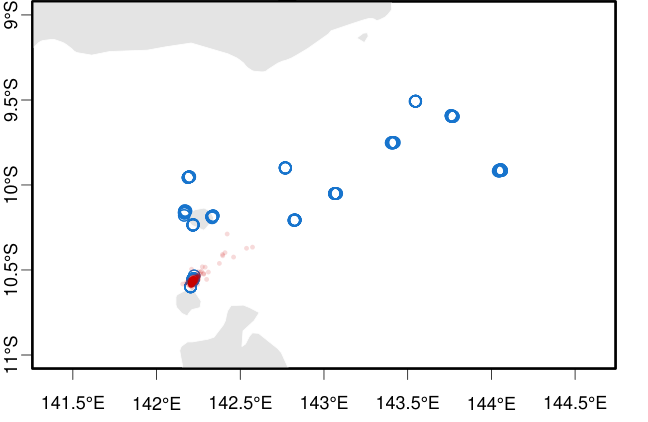

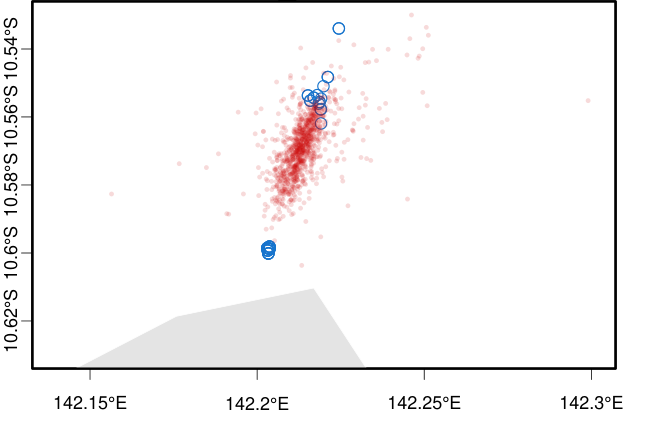

Figure S31. Origin tracing using Locator of sample TI23_044, collected on Waiben (Thursday Island) in 2023. Blue circles show reference samples locations. Red circles show n=1000 bootstrap replicates of inferred origin. Top plot shows all Torres Strait Islands, bottom plot is the same analysis showing details around Keriri (top blue cluster) and Muralug (bottom blue cluster).

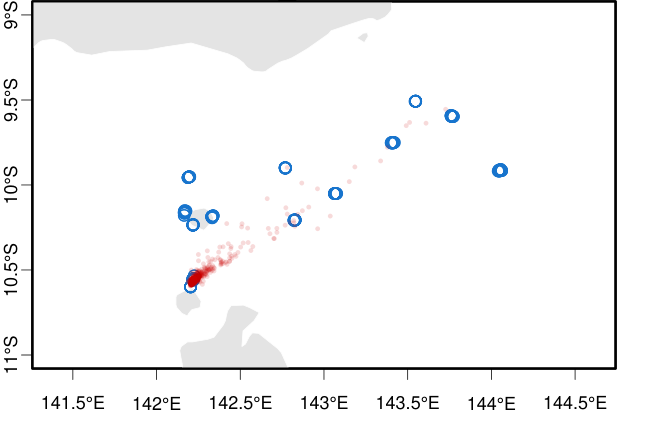

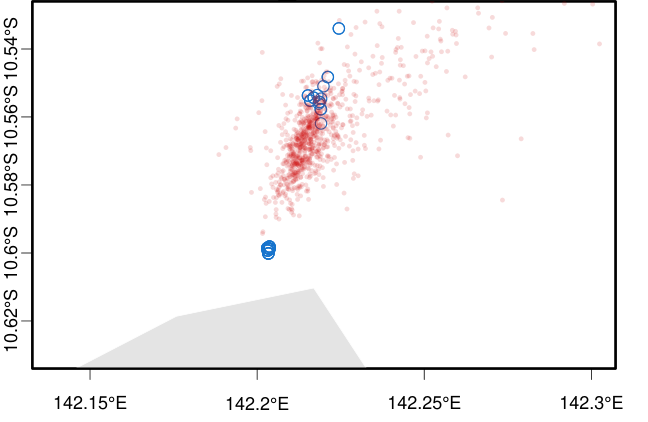

Figure S32. Origin tracing using Locator of sample TI23_045, collected on Waiben (Thursday Island) in 2023. Blue circles show reference samples locations. Red circles show n=1000 bootstrap replicates of inferred origin. Top plot shows all Torres Strait Islands, bottom plot is the same analysis showing details around Keriri (top blue cluster) and Muralug (bottom blue cluster).

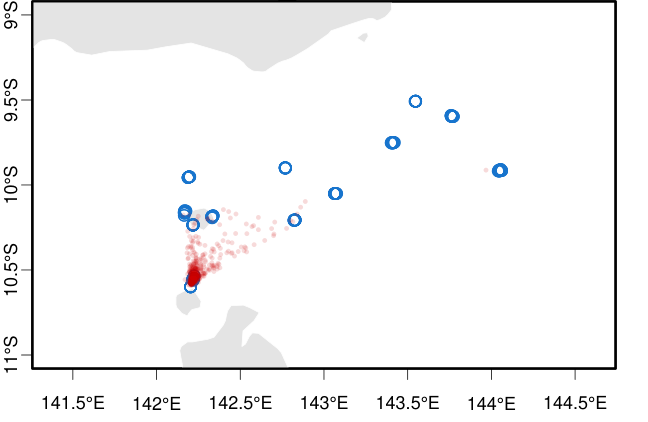

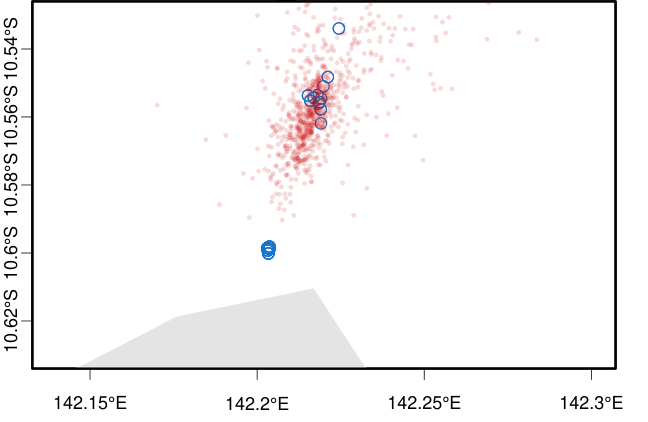

Figure S33. Origin tracing using Locator of sample TI23_046, collected on Waiben (Thursday Island) in 2023. Blue circles show reference samples locations. Red circles show n=1000 bootstrap replicates of inferred origin. Top plot shows all Torres Strait Islands, bottom plot is the same analysis showing details around Keriri (top blue cluster) and Muralug (bottom blue cluster).

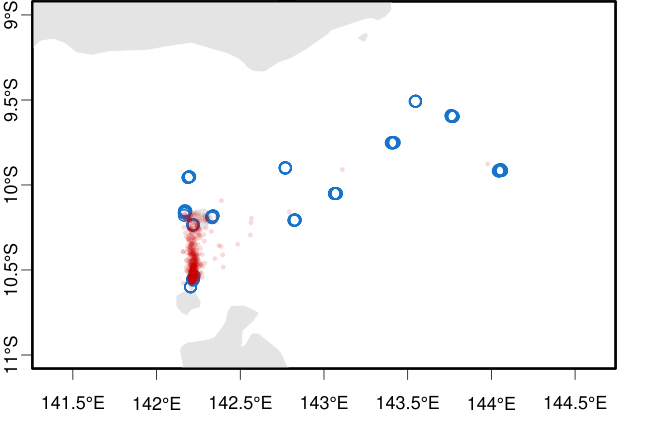

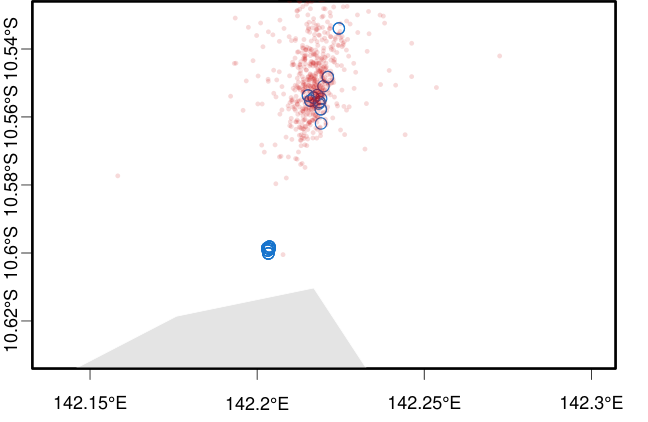

Figure S34. Origin tracing using Locator of sample TI23_047, collected on Waiben (Thursday Island) in 2023. Blue circles show reference samples locations. Red circles show n=1000 bootstrap replicates of inferred origin. Top plot shows all Torres Strait Islands, bottom plot is the same analysis showing details around Keriri (top blue cluster) and Muralug (bottom blue cluster).

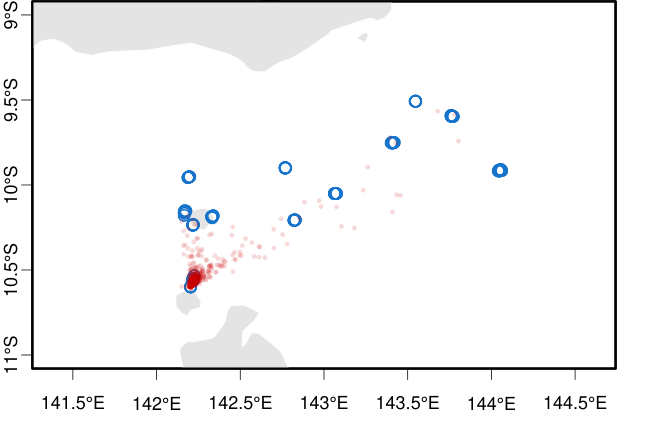

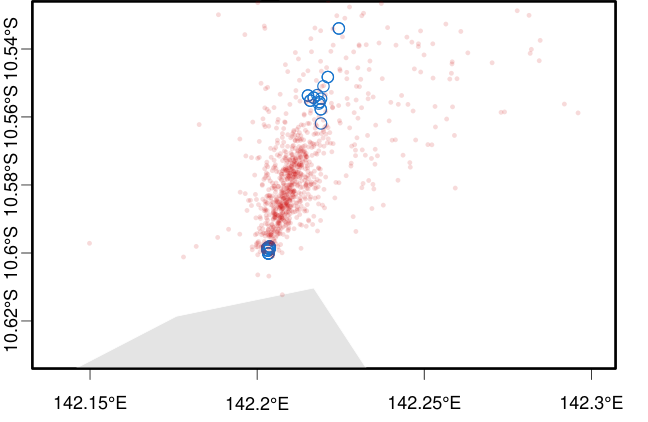

Figure S35. Origin tracing using Locator of sample TI23_048_1, collected on Waiben (Thursday Island) in 2023. Blue circles show reference samples locations. Red circles show n=1000 bootstrap replicates of inferred origin. Top plot shows all Torres Strait Islands, bottom plot is the same analysis showing details around Keriri (top blue cluster) and Muralug (bottom blue cluster).

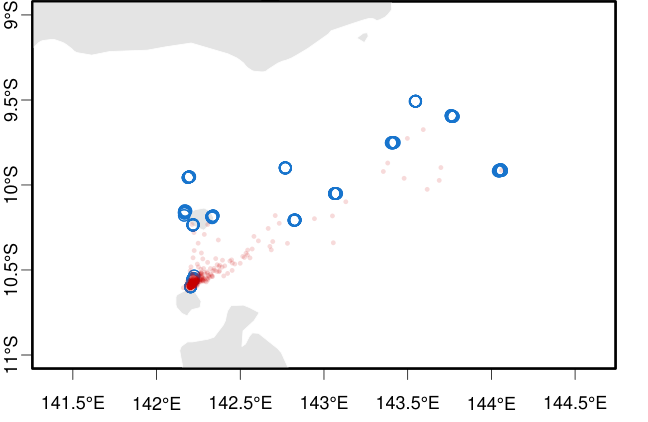

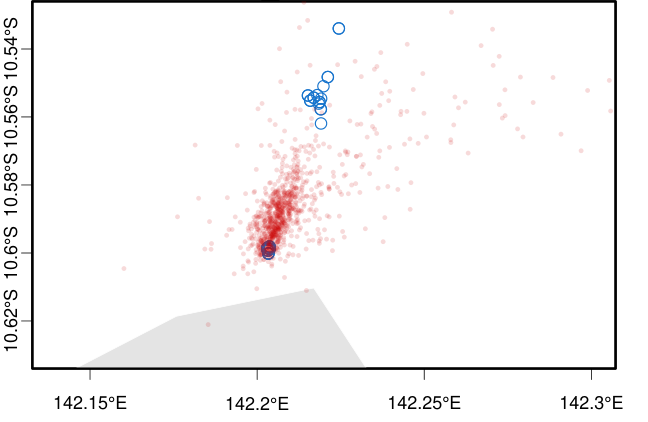

Figure S36. Origin tracing using Locator of sample TI23_048_2, collected on Waiben (Thursday Island) in 2023. Blue circles show reference samples locations. Red circles show n=1000 bootstrap replicates of inferred origin. Top plot shows all Torres Strait Islands, bottom plot is the same analysis showing details around Keriri (top blue cluster) and Muralug (bottom blue cluster).

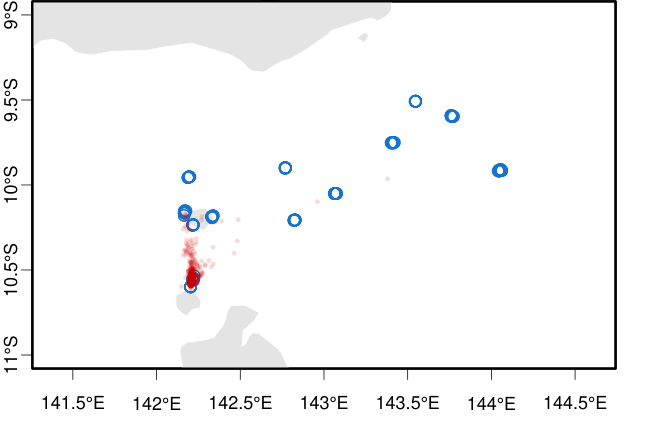

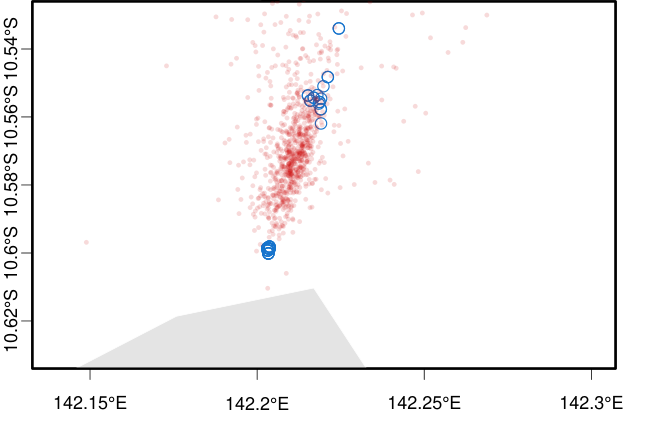

Figure S37. Origin tracing using Locator of sample TI23_048_3, collected on Waiben (Thursday Island) in 2023. Blue circles show reference samples locations. Red circles show n=1000 bootstrap replicates of inferred origin. Top plot shows all Torres Strait Islands, bottom plot is the same analysis showing details around Keriri (top blue cluster) and Muralug (bottom blue cluster).

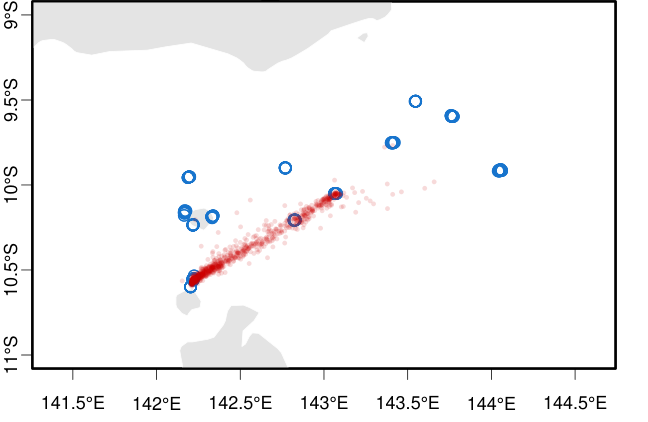

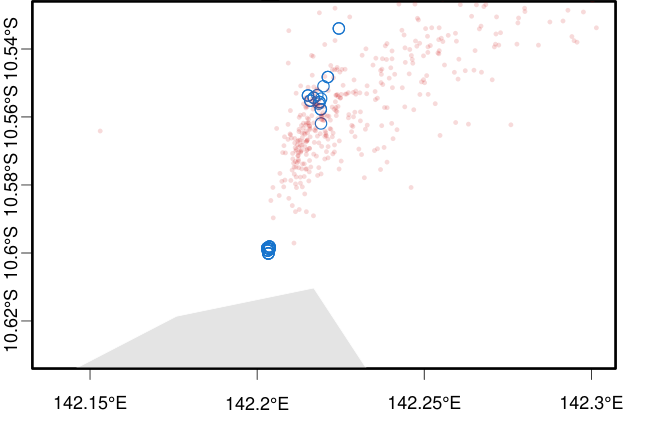

Figure S38. Origin tracing using Locator of sample TI23_048_4, collected on Waiben (Thursday Island) in 2023. Blue circles show reference samples locations. Red circles show n=1000 bootstrap replicates of inferred origin. Top plot shows all Torres Strait Islands, bottom plot is the same analysis showing details around Keriri (top blue cluster) and Muralug (bottom blue cluster).

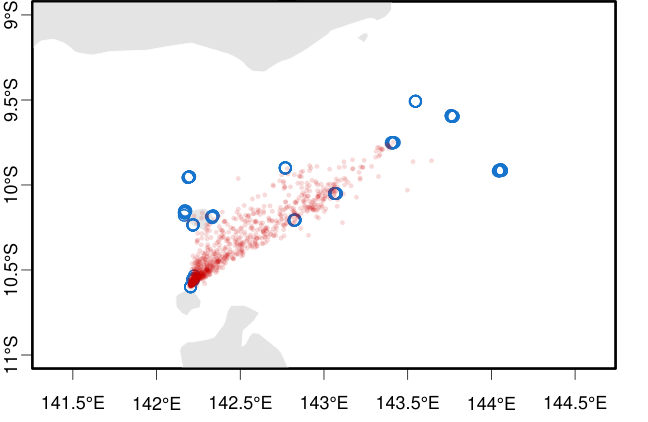

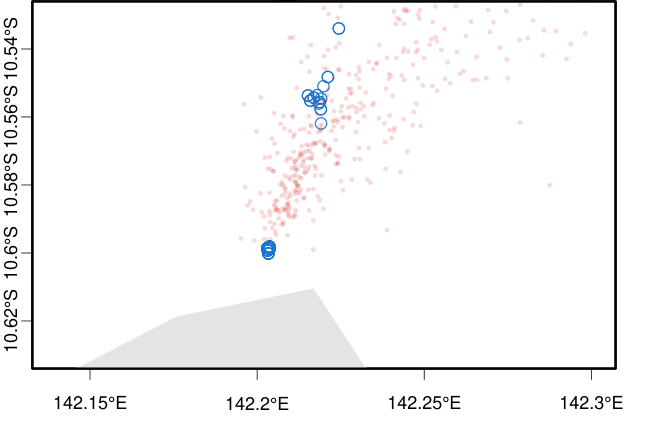

Figure S39. Origin tracing using Locator of sample TI23_049, collected on Waiben (Thursday Island) in 2023. Blue circles show reference samples locations. Red circles show n=1000 bootstrap replicates of inferred origin. Top plot shows all Torres Strait Islands, bottom plot is the same analysis showing details around Keriri (top blue cluster) and Muralug (bottom blue cluster).

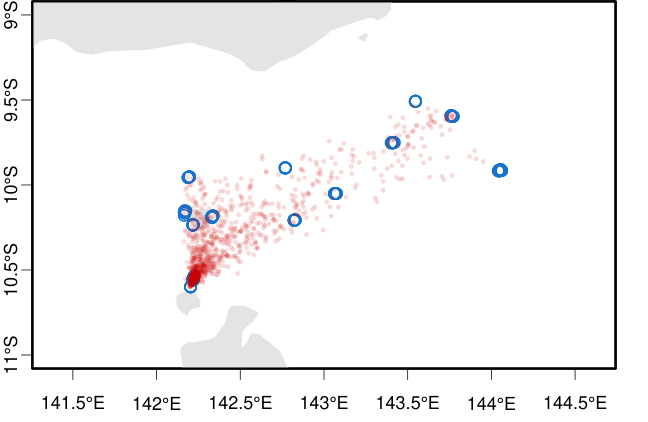

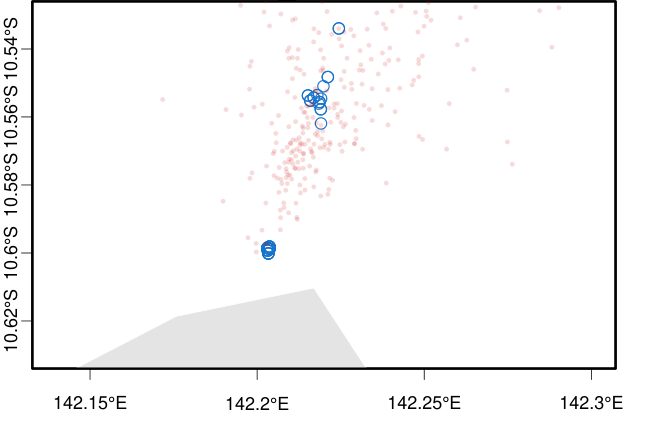

Figure S40. Origin tracing using Locator of sample TI23_050, collected on Waiben (Thursday Island) in 2023. Blue circles show reference samples locations. Red circles show n=1000 bootstrap replicates of inferred origin. Top plot shows all Torres Strait Islands, bottom plot is the same analysis showing details around Keriri (top blue cluster) and Muralug (bottom blue cluster).

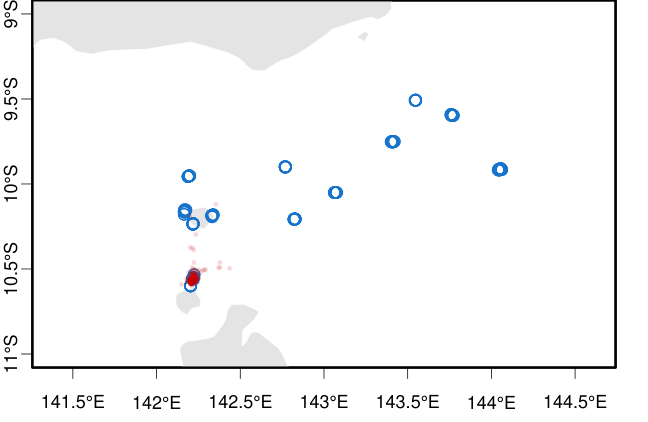

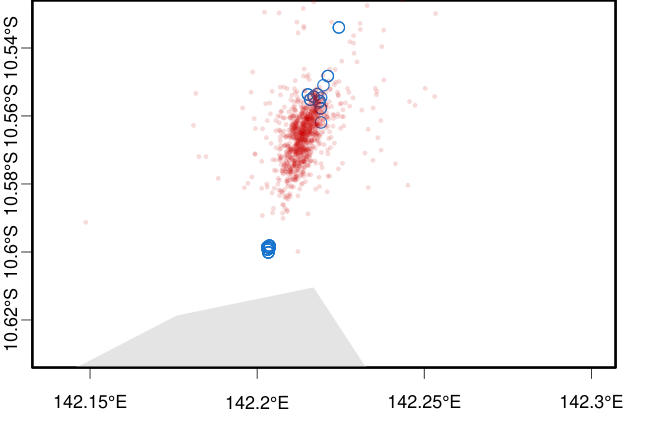

Figure S41. Origin tracing using Locator of sample TI23_051, collected on Waiben (Thursday Island) in 2023. Blue circles show reference samples locations. Red circles show n=1000 bootstrap replicates of inferred origin. Top plot shows all Torres Strait Islands, bottom plot is the same analysis showing details around Keriri (top blue cluster) and Muralug (bottom blue cluster).

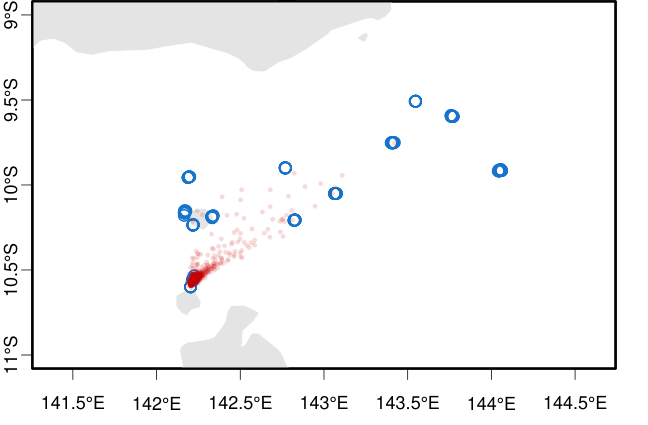

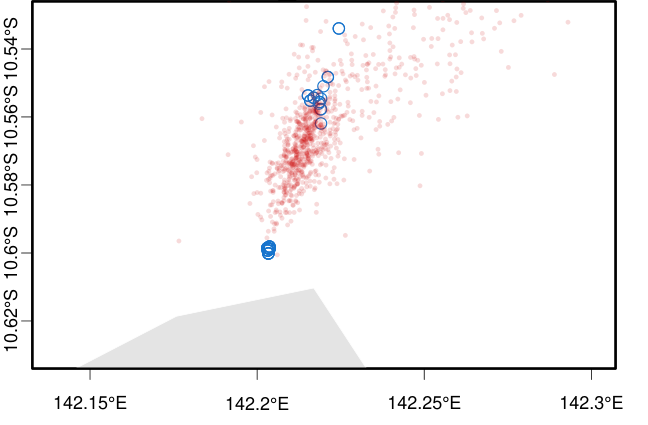

Figure S42. Origin tracing using Locator of sample TI23_052, collected on Waiben (Thursday Island) in 2023. Blue circles show reference samples locations. Red circles show n=1000 bootstrap replicates of inferred origin. Top plot shows all Torres Strait Islands, bottom plot is the same analysis showing details around Keriri (top blue cluster) and Muralug (bottom blue cluster).

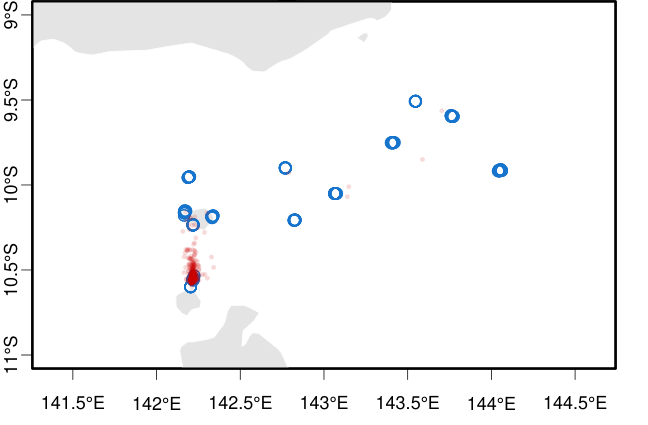

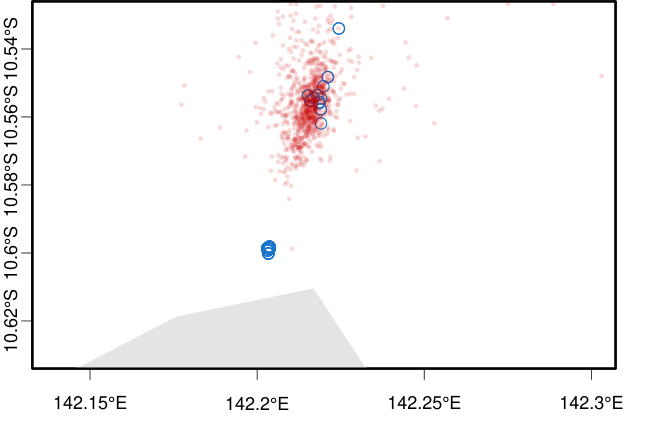

Figure S43. Origin tracing using Locator of sample TI23_053, collected on Waiben (Thursday Island) in 2023. Blue circles show reference samples locations. Red circles show n=1000 bootstrap replicates of inferred origin. Top plot shows all Torres Strait Islands, bottom plot is the same analysis showing details around Keriri (top blue cluster) and Muralug (bottom blue cluster).

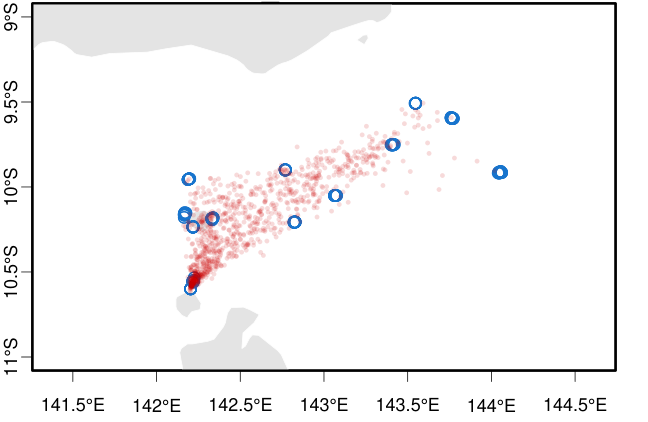

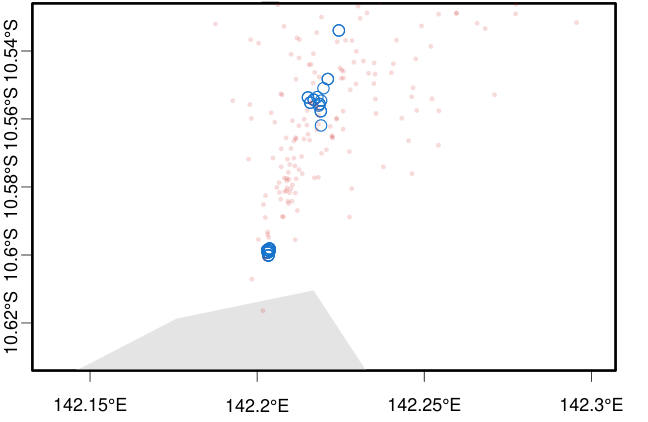

Figure S44. Origin tracing using Locator of sample TI23_054_1, collected on Waiben (Thursday Island) in 2023. Blue circles show reference samples locations. Red circles show n=1000 bootstrap replicates of inferred origin. Top plot shows all Torres Strait Islands, bottom plot is the same analysis showing details around Keriri (top blue cluster) and Muralug (bottom blue cluster).

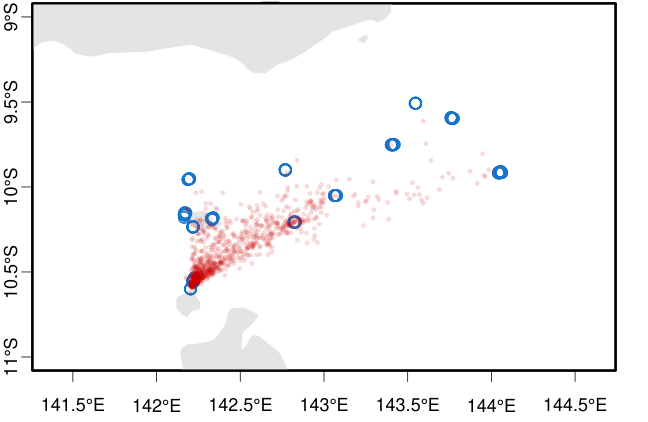

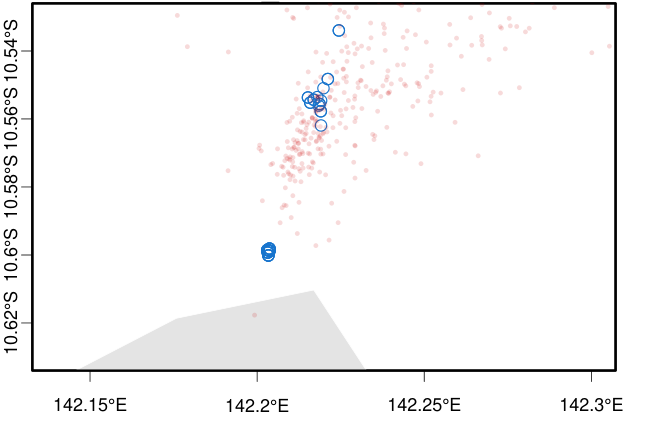

Figure S45. Origin tracing using Locator of sample TI23_054_2, collected on Waiben (Thursday Island) in 2023. Blue circles show reference samples locations. Red circles show n=1000 bootstrap replicates of inferred origin. Top plot shows all Torres Strait Islands, bottom plot is the same analysis showing details around Keriri (top blue cluster) and Muralug (bottom blue cluster).

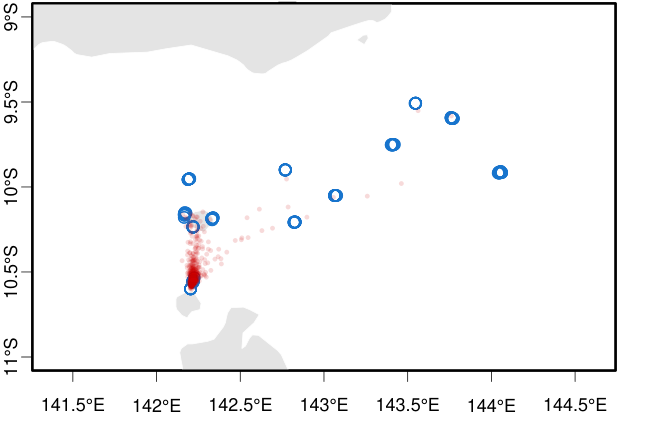

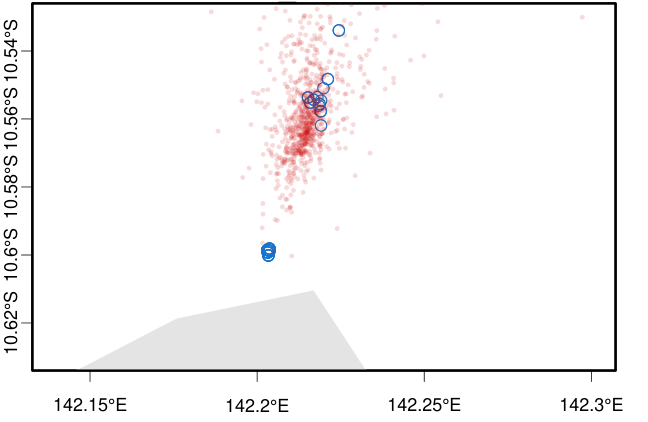

Figure S46. Origin tracing using Locator of sample TI23_055, collected on Waiben (Thursday Island) in 2023. Blue circles show reference samples locations. Red circles show n=1000 bootstrap replicates of inferred origin. Top plot shows all Torres Strait Islands, bottom plot is the same analysis showing details around Keriri (top blue cluster) and Muralug (bottom blue cluster).

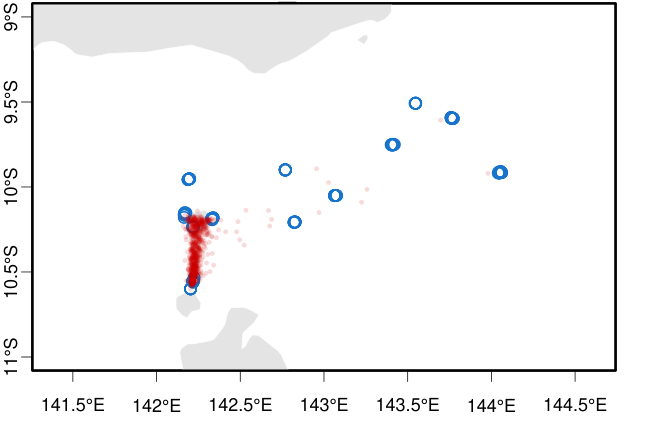

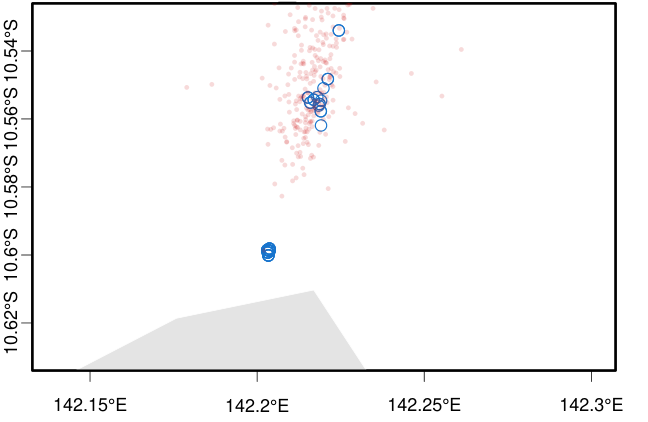

Figure S47. Origin tracing using Locator of sample TI23_056, collected on Waiben (Thursday Island) in 2023. Blue circles show reference samples locations. Red circles show n=1000 bootstrap replicates of inferred origin. Top plot shows all Torres Strait Islands, bottom plot is the same analysis showing details around Keriri (top blue cluster) and Muralug (bottom blue cluster).

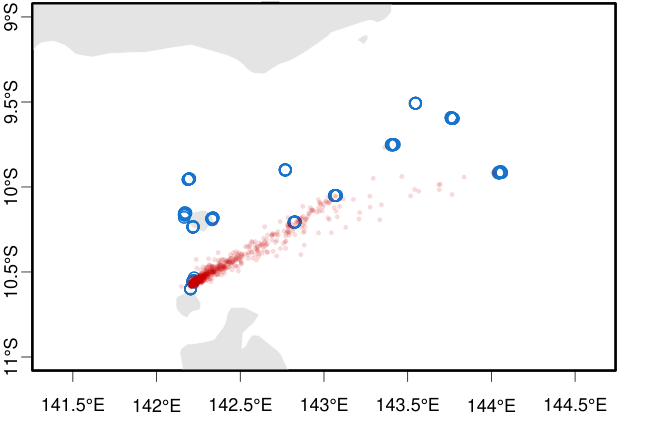

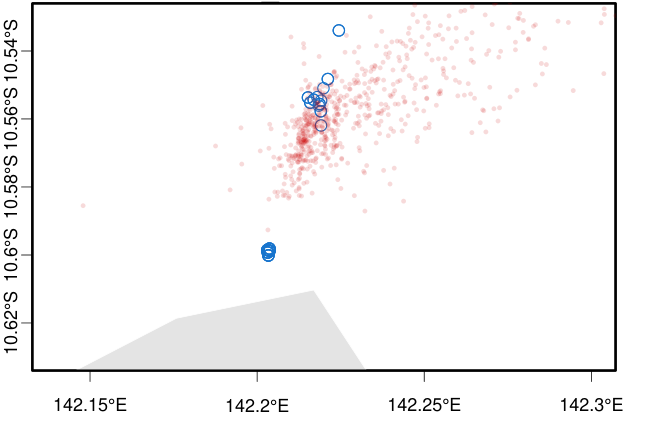

Figure S48. Origin tracing using Locator of sample TI23_057_1, collected on Waiben (Thursday Island) in 2023. Blue circles show reference samples locations. Red circles show n=1000 bootstrap replicates of inferred origin. Top plot shows all Torres Strait Islands, bottom plot is the same analysis showing details around Keriri (top blue cluster) and Muralug (bottom blue cluster).

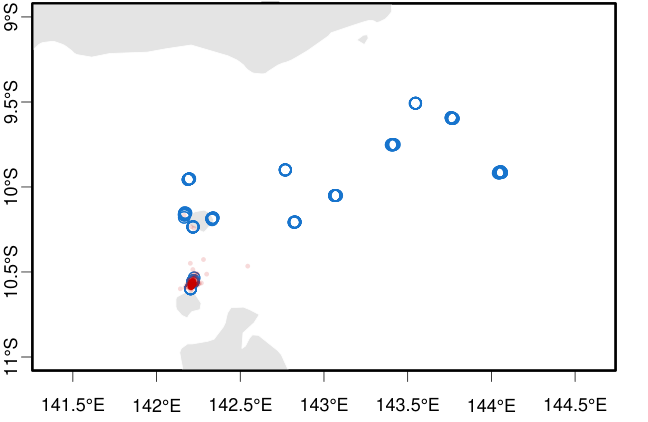

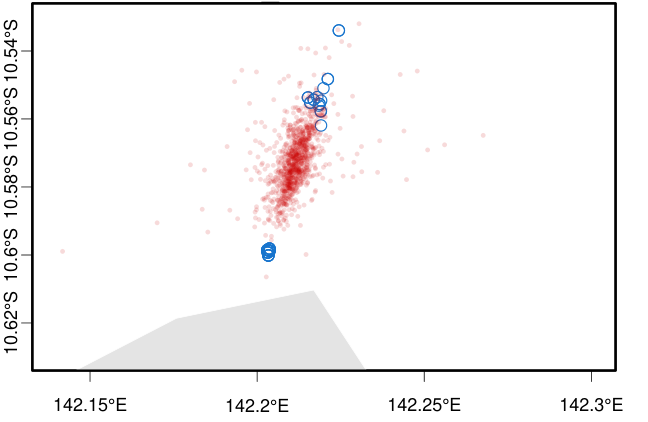

Figure S49. Origin tracing using Locator of sample TI23_057_2, collected on Waiben (Thursday Island) in 2023. Blue circles show reference samples locations. Red circles show n=1000 bootstrap replicates of inferred origin. Top plot shows all Torres Strait Islands, bottom plot is the same analysis showing details around Keriri (top blue cluster) and Muralug (bottom blue cluster).

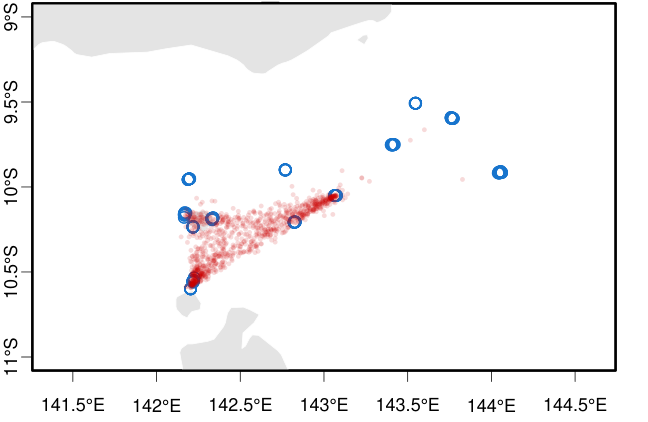

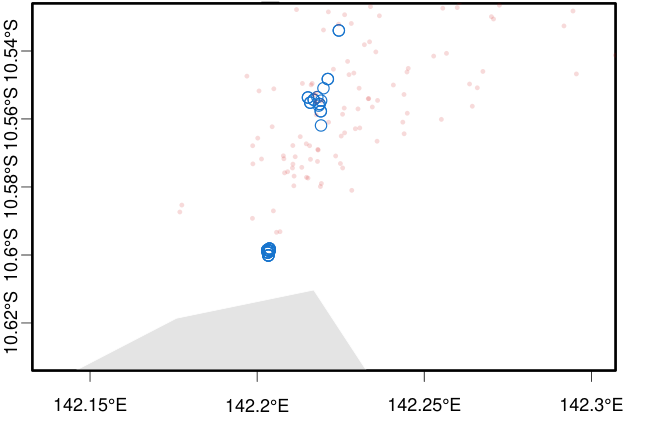

Figure S50. Origin tracing using Locator of sample TI23_057_3, collected on Waiben (Thursday Island) in 2023. Blue circles show reference samples locations. Red circles show n=1000 bootstrap replicates of inferred origin. Top plot shows all Torres Strait Islands, bottom plot is the same analysis showing details around Keriri (top blue cluster) and Muralug (bottom blue cluster).

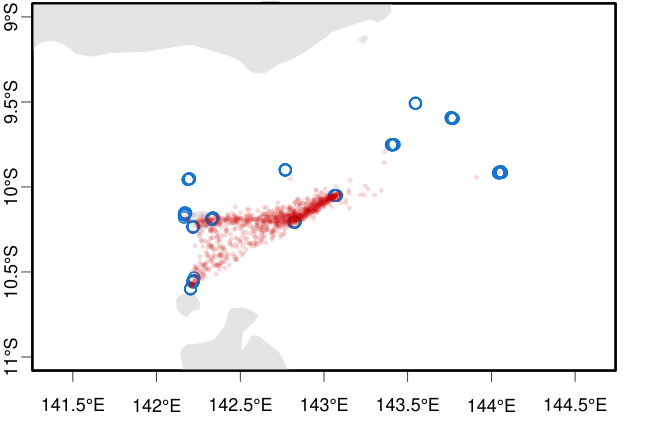

Figure S51. Origin tracing using Locator of sample TI23_057_4, collected on Waiben (Thursday Island) in 2023. Blue circles show reference samples locations. Red circles show n=1000 bootstrap replicates of inferred origin. Top plot shows all Torres Strait Islands, bottom plot is the same analysis showing details around Keriri (top blue cluster) and Muralug (bottom blue cluster).

Figure S52. Origin tracing using Locator of sample TI23_057_5, collected on Waiben (Thursday Island) in 2023. Blue circles show reference samples locations. Red circles show n=1000 bootstrap replicates of inferred origin. Top plot shows all Torres Strait Islands, bottom plot is the same analysis showing details around Keriri (top blue cluster) and Muralug (bottom blue cluster).

Figure S53. Origin tracing using Locator of sample TI23_058, collected on Waiben (Thursday Island) in 2023. Blue circles show reference samples locations. Red circles show n=1000 bootstrap replicates of inferred origin. Top plot shows all Torres Strait Islands, bottom plot is the same analysis showing details around Keriri (top blue cluster) and Muralug (bottom blue cluster).

Figure S54. Origin tracing using Locator of sample TI23_059, collected on Waiben (Thursday Island) in 2023. Blue circles show reference samples locations. Red circles show n=1000 bootstrap replicates of inferred origin. Top plot shows all Torres Strait Islands, bottom plot is the same analysis showing details around Keriri (top blue cluster) and Muralug (bottom blue cluster).

Figure S55. Origin tracing using Locator of sample TI23_060, collected on Waiben (Thursday Island) in 2023. Blue circles show reference samples locations. Red circles show n=1000 bootstrap replicates of inferred origin. Top plot shows all Torres Strait Islands, bottom plot is the same analysis showing details around Keriri (top blue cluster) and Muralug (bottom blue cluster).

Figure S56. Origin tracing using Locator of sample TI23_061, collected on Waiben (Thursday Island) in 2023. Blue circles show reference samples locations. Red circles show n=1000 bootstrap replicates of inferred origin. Top plot shows all Torres Strait Islands, bottom plot is the same analysis showing details around Keriri (top blue cluster) and Muralug (bottom blue cluster).

Figure S57. Origin tracing using Locator of sample TI25_014, collected on Waiben (Thursday Island) in 2025. Blue circles show reference samples locations. Red circles show n=1000 bootstrap replicates of inferred origin. Top plot shows all Torres Strait Islands, bottom plot is the same analysis showing details around Keriri (top blue cluster) and Muralug (bottom blue cluster).

Figure S58. Origin tracing using Locator of sample TI25_170, collected on Waiben (Thursday Island) in 2025. Blue circles show reference samples locations. Red circles show n=1000 bootstrap replicates of inferred origin. Top plot shows all Torres Strait Islands, bottom plot is the same analysis showing details around Keriri (top blue cluster) and Muralug (bottom blue cluster).

Figure S59. Origin tracing using Locator of sample TI25_188, collected on Waiben (Thursday Island) in 2025. Blue circles show reference samples locations. Red circles show n=1000 bootstrap replicates of inferred origin. Top plot shows all Torres Strait Islands, bottom plot is the same analysis showing details around Keriri (top blue cluster) and Muralug (bottom blue cluster).

Figure S60. Origin tracing using Locator of sample TI25_040, collected on Waiben (Thursday Island) in 2025. Blue circles show reference samples locations. Red circles show n=1000 bootstrap replicates of inferred origin. Top plot shows all Torres Strait Islands, bottom plot is the same analysis showing details around Keriri (top blue cluster) and Muralug (bottom blue cluster).

Figure S61. Origin tracing using Locator of sample TI25_041, collected on Waiben (Thursday Island) in 2025. Blue circles show reference samples locations. Red circles show n=1000 bootstrap replicates of inferred origin. Top plot shows all Torres Strait Islands, bottom plot is the same analysis showing details around Keriri (top blue cluster) and Muralug (bottom blue cluster).

Figure S62. Origin tracing using Locator of sample TI25_059, collected on Waiben (Thursday Island) in 2025. Blue circles show reference samples locations. Red circles show n=1000 bootstrap replicates of inferred origin. Top plot shows all Torres Strait Islands, bottom plot is the same analysis showing details around Keriri (top blue cluster) and Muralug (bottom blue cluster).

Figure S63. Origin tracing using Locator of sample TI25_079, collected on Waiben (Thursday Island) in 2025. Blue circles show reference samples locations. Red circles show n=1000 bootstrap replicates of inferred origin. Top plot shows all Torres Strait Islands, bottom plot is the same analysis showing details around Keriri (top blue cluster) and Muralug (bottom blue cluster).

Figure S64. Origin tracing using Locator of sample TI26_015, collected on Waiben (Thursday Island) in 2026. Blue circles show reference samples locations. Red circles show n=1000 bootstrap replicates of inferred origin. Top plot shows all Torres Strait Islands, bottom plot is the same analysis showing details around Keriri (top blue cluster) and Muralug (bottom blue cluster).
